# A Functional Genetic Atlas of Parkin Resolves Variants of Uncertain Significance and Predicts Parkinson’s Disease Age at Onset

**DOI:** 10.64898/2026.02.09.704817

**Authors:** Vahid Aslanzadeh, Sophie Glendinning, Yustika Sari, Marguerite Clarke, Christine J. Rodger, Xiaoyi Lin, Dipti Ranjan Lenka, Lucy Richardson, Jin Rui Liang, Teresa Kleinz, Duncan Sproul, Katja Lohmann, Christine Klein, Esther M. Sammler, Dario R. Alessi, Atul Kumar, Miratul M.K. Muqit, Grzegorz Kudla

**Author notes:** These authors contributed equally to this study (Vahid Aslanzadeh, Sophie Glendinning, Yustika Sari). Joint corresponding authors.

## Abstract

Autosomal recessive mutations in the Parkin gene (*PRKN*) cause early-onset Parkinson’s disease (PD). Parkin functions as a ubiquitin E3 ligase acting downstream of the PINK1 kinase to promote phosphorylated ubiquitin at sites of mitochondrial damage. Yet the functional effects of most *PRKN* gene variants remain unknown. Here we use a pooled cellular assay measuring phosphorylated ubiquitin accumulation to quantify the activity of ∼9,200 *PRKN* missense and nonsense variants. Our screen identifies thousands of loss- and gain-of-function variants, including activating variants mainly residing at autoinhibitory interfaces. Functional scores accurately distinguish known PD pathogenic and benign variants, and enable reclassification of 173/184 variants of uncertain significance (VUS). When combined into biallelic genotype scores, these data predict the age at disease onset in patients, revealing a quantitative link between Parkin activity and clinical manifestation. Our results provide a comprehensive functional genetic map of Parkin and demonstrate the power of multiplexed assays of variant effects (MAVEs) for variant interpretation and precision medicine in PD.

## Introduction

Parkinson’s disease (PD) is a neurodegenerative disorder caused by the loss of dopaminergic neurons in the midbrain’s substantia nigra pars compacta, often associated with the formation of cytoplasmic aggregates known as Lewy bodies (LB) [1]. It affects approximately 1% of people over the age of 60 years, and is characterised by motor and non-motor symptoms including bradykinesia, tremor, rigidity, cognitive decline, and mood disorders. While current treatments can alleviate symptoms, no therapy has been proven to halt or reverse disease progression [2]. The aetiology of PD is not fully understood, although a combination of ageing, environmental factors, and genetics play a role in its pathogenesis [3, 4]. About 15% of PD patients have a known genetic cause or strong genetic contribution, most commonly in genes associated with endolysosomal pathways (*LRRK2*, *VPS35, GBA1*), mitochondria quality control (*PINK1*, *PRKN*, *PARK7*) or protein misfolding and LB formation (*SNCA*) [5, 6].

Autosomal recessive variants in the *PRKN* (*PARK2*) gene account for <1% of all cases, but the majority of early-onset PD, including 77% of juvenile PD patients [7]. *PRKN* encodes Parkin, a ubiquitin E3 ligase that together with the kinase PINK1 regulates the ubiquitin-dependent clearance of damaged mitochondria by autophagy (mitophagy) [8, 9]. Parkin belongs to the RING-between-RING (RBR) family of E3 ligases, and contains an N-terminal ubiquitin-like (UBL) domain, activating element (ACT) and RING0 domains, followed by RING1, in-between-RING (IBR), repressor element (REP) and RING2 domains that mediate the transfer of ubiquitin from an E2 ubiquitin-conjugating enzyme to target proteins. In its native state, Parkin is autoinhibited, via three major interfaces: the E2-binding site on RING1 is occluded by REP and UBL domains; the donor Ub binding site on RING1 by UBL-IBR interface; and the catalytic C431 residue in RING2 by RING0 [10–12]. Activation of Parkin is triggered by initial PINK1-dependent phosphorylation of ubiquitin at Serine 65 (pSer65-Ub) [13–15]. pSer65-Ub binds with high affinity to Parkin and recruits it to sites of damaged mitochondria. This leads to a conformational change that allows phosphorylation of S65 in the Parkin UBL domain by PINK1 [16], leading to further conformational changes that allow catalytic C431 to be transthiolated by E2-Ub [10–12]. Parkin activation, ubiquitylation of substrates and subsequent phosphorylation by PINK1 is rapid due to feed-forward amplification and as such pSer65-Ub accumulation can be used a measure of Parkin activity in cells [17, 18].

Hundreds of *PRKN* variants have been observed in PD patients (https://www.mdsgene.org/), but many remain classified as variants of uncertain significance (VUS) and thousands more are likely to exist in the general population but have not yet been catalogued. The majority of juvenile and early-onset PD patients harbour biallelic variants, while heterozygous variants have been linked to late-onset PD risk [19, 20]. The mutational spectrum is broad, and pathogenic variants spread over the entire gene, including truncating and missense as well as copy number variants [21, 22]. Many of the PD missense variants affect residues critical for Parkin regulation or catalytic activity, including S65, K161, K211 and C431, but most have not been functionally assessed. Prior biochemical and structural studies of Parkin have uncovered several activating mutations that have stimulated drug development strategies [23–27], however, their clinical relevance is unclear. A recent saturation mutagenesis study mapped effects on protein abundance [28], but this captured only a subset of *PRKN* pathogenic variants that decreased abundance, highlighting the need for more studies of catalytic function. With increasing clinical exome and whole genome sequencing of PD patients in the clinic, the ability to mechanistically interpret genetic data will be of high value in aiding genetic counselling of PD patients and informing trials of disease modifying therapies currently underway for mitophagy-enhancing therapies [9]. Further, this knowledge will be valuable globally as more genetic data from non-Caucasian populations emerges including from the Global Parkinson’s Genetics Program (GP2) [29].

Here we describe a multiplex assay of variant effects (MAVE) of *PRKN*, using cellular accumulation of phosphorylated ubiquitin (pSer65-Ub), following mitochondrial depolarisation-induced damage, as a readout of Parkin activity. Robustness of the assay was validated by observation of complete loss of Parkin activity for variants affecting all known critical regulatory amino acids including the activating phosphorylation residue, S65, and the catalytic residue, C431. The assay accurately classifies nearly all known benign and pathogenic variants, providing functional evidence for 173 of 184 variants of uncertain significance (VUS), to help reclassify them as either benign or pathogenic. When combined with abundance data and used to calculate biallelic pathogenicity scores, our data reveal a robust relationship between Parkin mutation status and age at disease onset.

## Results

### Design and validation of Parkin activity screen

To enable the mutational screen, we designed a fluorescence activated cell sorting (FACS)-based assay of phosphorylated ubiquitin (pSer65-Ub) in HeLa cells as a functional readout of Parkin catalytic activity. Upon mitochondrial depolarisation, PINK1-mediated activation of Parkin induces pSer65-Ub accumulation on the outer mitochondrial membrane [8, 9]; in the absence of Parkin, the pSer65-Ub response to mitochondrial damage is significantly reduced. We replicated these results in HeLa cells expressing single-copy Parkin alleles from a genomic landing pad. Following mitochondrial depolarisation induced by 1 μM Oligomycin and 10 μM Antimycin A (OA), we observed time-dependent pSer65-Ub accumulation in cells expressing WT Parkin but not in those expressing the catalytically inactive C431F or the phosphorylation-deficient S65A mutant (**Fig. 1A,B**), validating the assay. In cells expressing these inactive mutants, OA treatment led to only a modest increase in pSer65-Ub at later timepoints (60-80 min), due to Parkin-independent mechanisms via as yet unknown ubiquitin E3 ligases (**Fig 1B**). Using the panel of functional and nonfunctional Parkin variants, we then optimised the FACS assay, selecting 45 minutes OA treatment, as this corresponded to the linear phase of the WT Parkin enzymatic reaction and yielded the best dynamic range for Parkin activity detection (**Fig 1B**).

**Figure 1.**
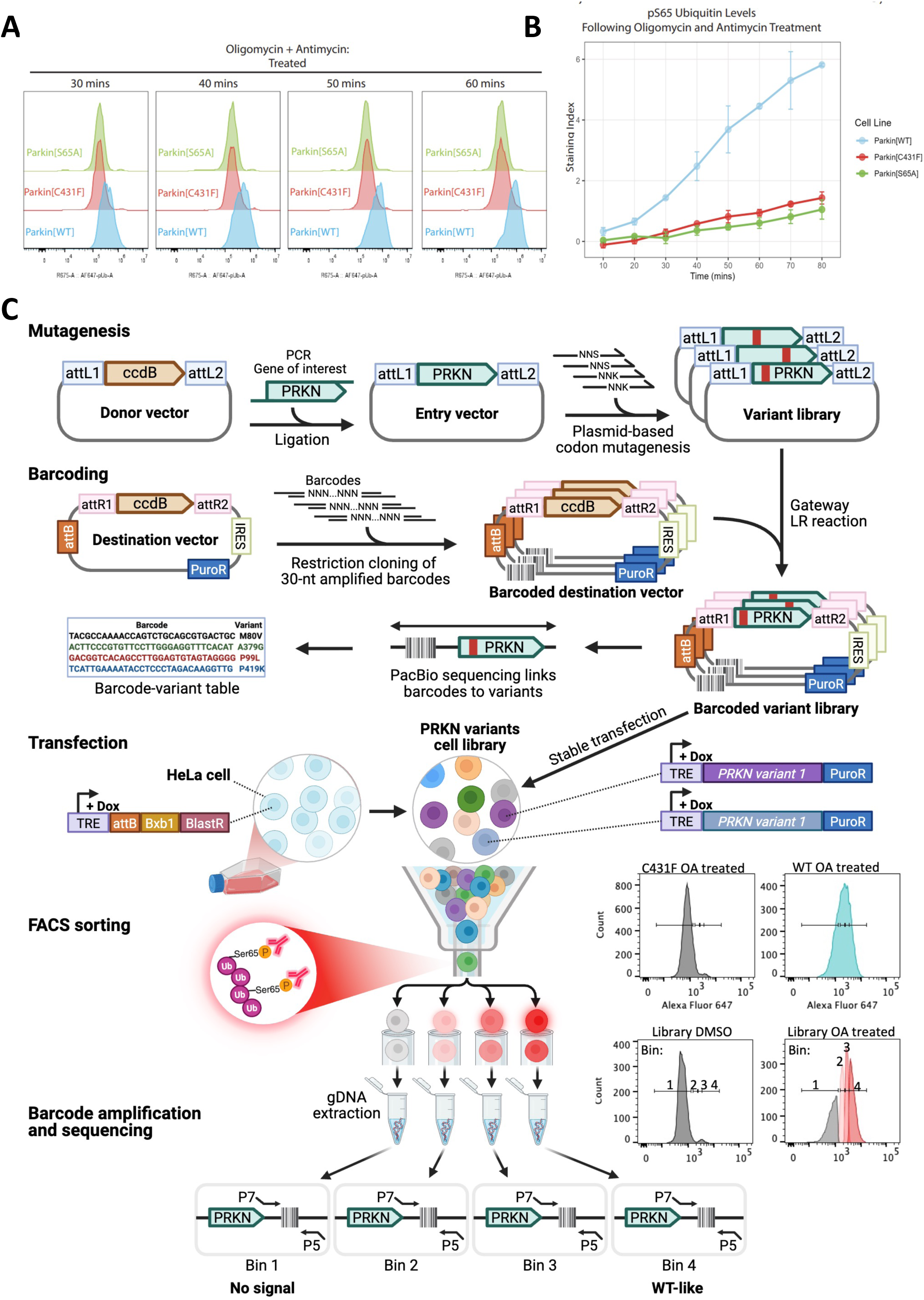
Development of Parkin MAVE platform. **A)** Time course of pS65 ubiquitin levels in HeLa BxB1 cells expressing Parkin[WT], Parkin[C431F], or Parkin[S65A], measured by flow cytometry within the indicated time after 1 uM Oligomycin and 10 uM Antimycin treatment. **B)** Time course of pS65-Ub evaluated using the staining index, the geometric mean fluorescence intensity normalized to vehicle control / robust SD of vehicle control. Points represent mean ± SD from two independent experiments. **C)** Schematic of Parkin MAVE platform, indicating, from top to bottom: saturation mutagenesis of Parkin; generation and sequencing of barcoded variant library; pooled transfection into Hela with a doxycycline-inducible Bxb1-targetable landing pad; flow sorting by pS65-Ub intensity; extraction of genomic DNA and generation of barcode libraries for Illumina sequencing. Figure created in BioRender.

We then cloned the *PRKN* gene into pAINt [30], a Gateway entry vector compatible with a published nicking mutagenesis protocol [31], and used saturation mutagenesis to generate 32 variants per codon, starting from the second codon (**Fig. 1C**). In parallel, we integrated a library of 20 million barcodes into the pMV2 destination vector backbone. The *PRKN* mutant library was then recombined into barcoded pMV2 via a Gateway LR reaction to generate 9,234 single missense and stop codon variants out of 9,280 possible (99.5% coverage), tagged by 438,157 unique barcodes. 99% of variants were associated with 5 barcodes or more; 24,235 barcodes were associated with synonymous variants and 146,793 barcodes were linked with unmutated *PRKN* and were taken as WT in downstream analyses (**Supplementary Table 1, Supplementary Fig. 1A-D**). The barcoded Parkin plasmid library was introduced into HeLa-Bxb1 landing pad cells by electroporation and selected with puromycin for two weeks to keep single integrants (**Fig. 1C**). To assess Parkin function, cells were treated with OA for 45 minutes, permeabilized, and probed with pSer65-Ub primary antibody, followed by an AF647-labeled secondary antibody (**Fig. 1C**). Cells were subsequently FACS-sorted into four bins based on AF647 fluorescence intensity. From every bin DNA was extracted, barcodes were amplified, and high-throughput sequencing was used to quantify barcode frequency per bin (**Fig. 1C**). After removing barcodes with fewer than 150 reads across 4 bins, a weighted average was calculated for each barcode based on its distribution across bins, and a score calculated for each variant. False discovery rates were computed by a bootstrapping approach using barcode scores per variant in the library.

### Functional landscape of Parkin variants

We obtained functional scores for 9,176 (98.8%) missense and nonsense *PRKN* variants, with an average of 19 barcodes per variant (**Fig 2A, Supplementary Fig. 1B, Supplementary Table 2**). The average score of synonymous variants was indistinguishable from WT (WT score = 0, synonymous score=-0.016, t-test p-value > 0.8), whereas almost all nonsense variants scored low (Stop score=-0.99), with the exception of a few scattered variants with low barcode coverage (which we interpret as experimental noise), and several stop codons near the C-terminus, indicating that the last few amino acids are not essential for function. The distribution of missense variant scores was bimodal: 75% were statistically indistinguishable from wild-type (FDR > 0.05 vs WT), while 13% were indistinguishable from complete loss-of-function variants (FDR > 0.05 vs stop codons), suggesting they retain full or no activity, respectively (**Fig. 2B**). The bimodal distribution likely reflects the ultrasensitive nature of the PINK1–Parkin feed-forward amplification loop, which amplifies small differences in Parkin activation into all-or-none outcomes. In addition, around 10% of missense variants only showed a partial loss of function and 2.5% (FDR < 0.05) were activating. The complete dataset can be accessed through our web app (https://parkin-app-umfr.onrender.com/) and MAVEDB.

**Figure 2.**
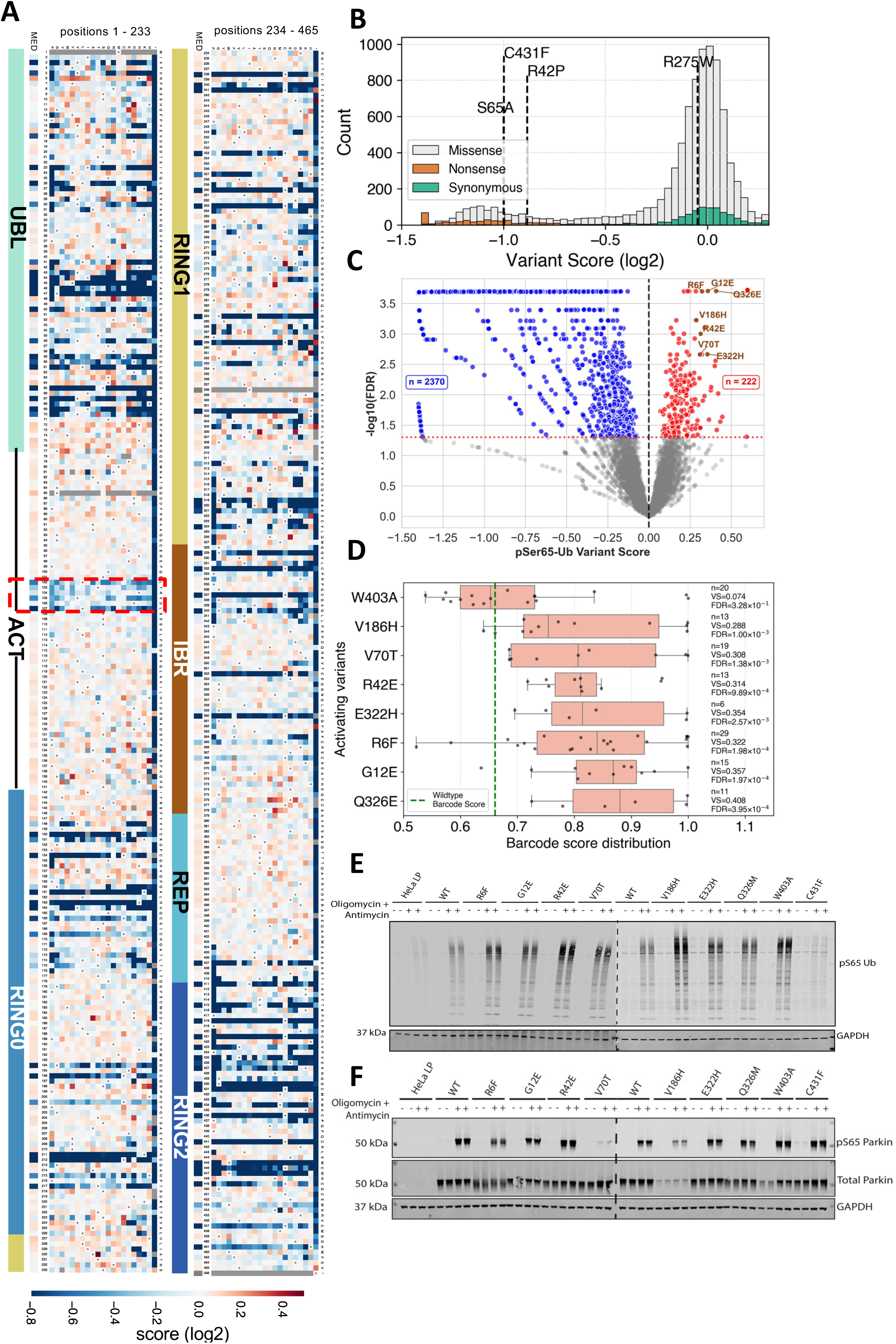
Functional map of Parkin variants. **A)** Functional scores of Parkin variants, showing levels of pS65-Ub measured 45 minutes after OA treatment, are indicated with a colour scale, with inactivating variants in blue, wild type-like variants in white, and activating variants in red. Variants with missing data are in grey. The domain architecture of Parkin is shown on the left. MED, median variant scores per position. **B)** Distribution of Parkin functional scores for all missense (grey), synonymous (green) and nonsense (orange) variants. The median score of wild type barcodes is set as 0 and the median score of nonsense variants is -1. **C)** Volcano plot of Parkin variants showing pS65-Ub scores and associated false discovery rates (vs wild-type Parkin). **D)** Distribution of barcode scores for candidate activating variants. **E-F)** Representative immunoblot analysis of pS65 Ubiquitin (E), and pS65 Parkin and total Parkin levels (F) within HeLa BxB1 untransfected cells and HeLa BxB1 cells stably expressing different Parkin variants. Cells were treated with 1 μM Oligomycin and 10 μM Antimycin for 45 minutes before lysis.

Several well-studied disease/functional variants were in the complete loss-of-function (LOF) category, including S65A, C431F, and R42P (**Fig. 2B**). The status of S65 variants is consistent with the role of S65 as the critical activating phosphorylation site mediated by PINK1. Almost all alterations in this position resulted in LOF, except S65T that retained near wild-type activity, indicating that PINK1 can phosphorylate threonine as well as serine at this position (**Supplementary Fig. 1E**). None of the variants of S108 were LOF (**Fig 2A)** which may be due to this phosphorylation event being maximal within 2-5 mins of OA stimulation and its functional role being less important at 45 min OA stimulation under which our screen was performed [32]. R275W is amongst the most frequent pathogenic variants of Parkin [22, 33, 34]. In our data, most R275 mutations had relatively mild effects, and R275W did not differ significantly from wild-type (FDR > 0.05), suggesting that the impact of R275W may be context-dependent and not fully captured in our assay conditions. We also observed partial loss of function caused by substitutions within the ACT domain (residues 102–107) (**Fig 2A**, dashed red box). This observation aligns with structural studies showing that ACT-mediated interactions at the hydrophobic patch of the RING0 domain facilitate RING2 domain release and subsequent Parkin activation [35–37].

Activating variants in Parkin have been previously described under recombinant in vitro studies and cell-based studies of over-expressed Parkin [23–27, 38]. We detected activating variants throughout the gene, with several positions appearing as hotspots; examples include R6, R42, R51, and R72. At R6, 10 out of 19 amino acid substitutions showed significantly higher activity than wild-type (FDR < 0.05) (**Fig 2A**). We prioritised seven high-confidence activating variants located across different structural domains (R6F, G12E, R42E, V70T, V186H, E322H, Q326E) for follow-up experiments and also included W403A, which has been previously reported as activating (**Fig. 2C,D**) [24–26]. To determine the effect of the selected Parkin variants on activation, HeLa cell lines stably expressing these variants were treated with or without OA for 45 mins to induce mitochondrial depolarisation. Immunoblot analysis of whole-cell extracts confirmed that all variants with the possible exception of E322H caused an increase of pSer65-Ub relative to WT Parkin following OA treatment (**Fig. 2E, Supplementary Fig. 2**; median increase 1.5-fold). As expected, control cells without Parkin or those with the C431F variant showed negligible pSer65-Ub signals. Increased pSer65-Ub amongst variants was not accompanied by an increase of Parkin abundance, but some activating variants showed decreased Parkin levels (most notably V186H), indicating destabilisation-associated activation. Other variants exhibited an increased ratio of phosphorylated to total Parkin (W403A), or a slightly altered migration pattern (R6F) (**Fig. 2F**), suggesting that changes in post-translational regulation rather than increased Parkin levels underlie the increased activity.

### Clinical classification of variants and benchmarking of predictors

To test the utility of our data for resolving variants of unknown significance (VUS), we first compared functional scores with clinical annotations of *PRKN* variants. We used two publicly available databases, MDSGene (https://www.mdsgene.org/) and ClinVar (https://www.ncbi.nlm.nih.gov/clinvar/), that both report genetic variants in relation to phenotypic presentation in patients. They also assign a pathogenicity scoring to each variant that is based on its frequency in patients and controls, in-silico prediction and, if available, on experimental evidence. MDSGene includes carefully curated data from the published literature, and ClinVar is populated by diagnostic laboratories and researchers. We herein used both resources and merged the MDSGene and ClinVar datasets, prioritising MDSGene classifications and supplementing with ClinVar when a variant was unclassified or missing from MDSGene (e.g., A82E). Among variants annotated as benign or likely benign (n = 15), 14 had wild-type-like scores in our assay. Conversely, 21 of 24 pathogenic (P) or likely pathogenic (LP) variants showed strong loss of function (**Fig. 3A**). Three P/LP variants—R33Q, R275W, and G328E—retained wild-type-like activity despite evidence of pathogenicity in patients, for instance, by large families with segregation and recurrence. Biochemical and cell-based studies of these variants previously returned ambiguous results, e.g. the G328E mutant which has been identified in early-onset PD [7] is located within the IBR domain and does not structurally affect the protein fold [39]. Furthermore, its effect based on *in vitro* assays has been conflicting with reports of reduced activity [24] or slightly enhanced activity [40]. Similarly, the R33Q mutant within the UBL domain exhibits increased Parkin activity by *in vitro* assays [23, 40]. Structurally, the effects of G328E and R33Q are less clear and they may affect interdomain interactions or interaction/binding with other proteins [41] which may not be reflected by measuring accumulation of pSer65-Ub. Among VUS, most showed wild-type-like activity (**Fig. 3A**), consistent with the overall distribution of functional effects in our dataset, however, a small subset showed complete loss of function, including S65N, a residue essential for phosphorylation-dependent Parkin activation, and C154R and C337Y, both of which are within Zn-binding motifs essential for the structural stability of Parkin. Our previous analysis of Parkin in primary fibroblasts from a homozygous S65N PD patient demonstrated complete loss of activity [42]. However, C154R and C337Y have been reported in single individuals only and S65N was reported in two unrelated individuals [42], resulting in unclassified status. These results underline the importance of functional tests to distinguish pathogenic and benign variants and our results provide a strong indication that these variants are, in fact, pathogenic.

**Figure 3.**
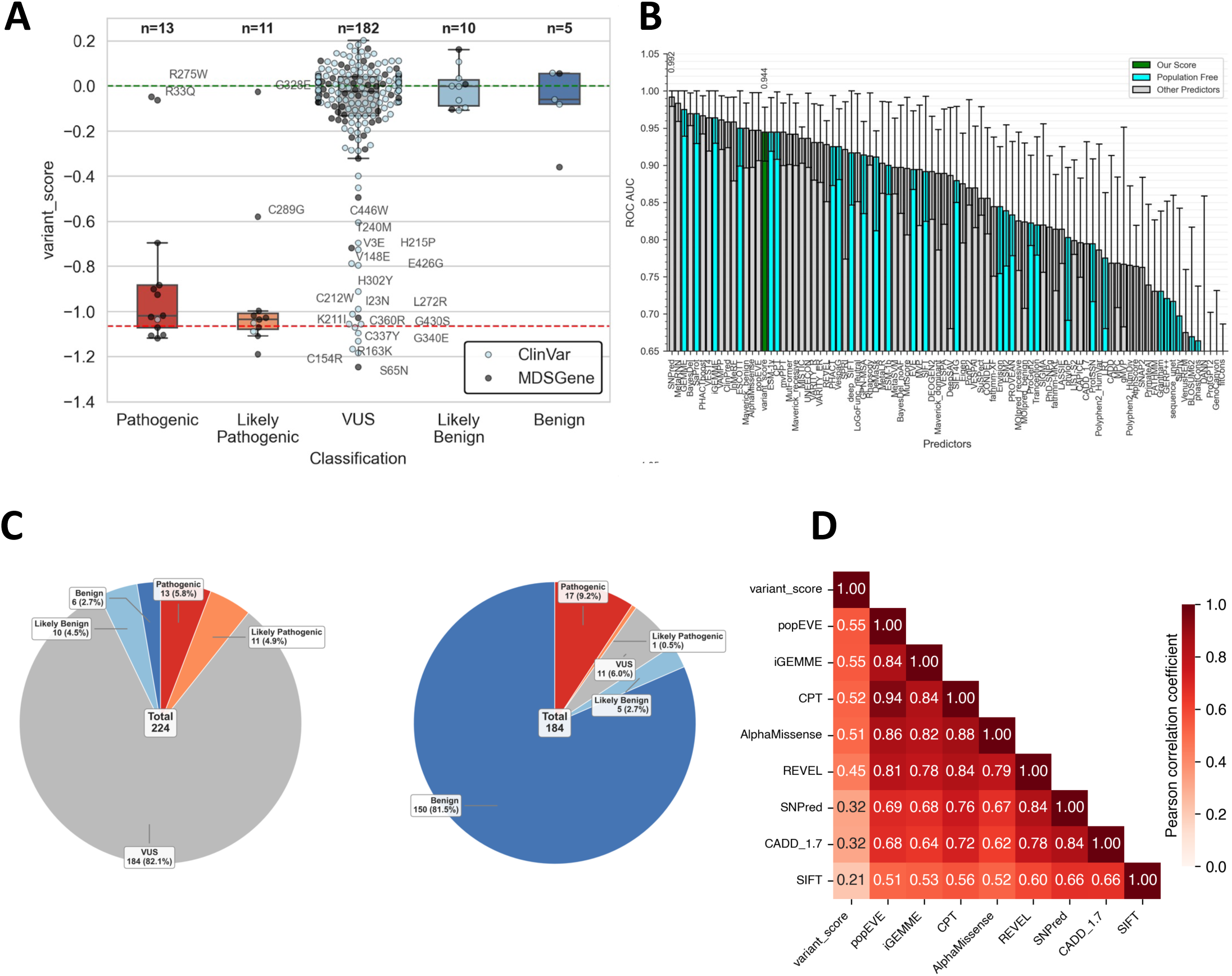
Clinical variant interpretation. **A)** Boxplots showing pS65-Ub functional scores for variants stratified according to their clinical annotations. Boxes show the interquartile range (IQR) with median line, whiskers denote 1.5 × IQR. MDSGene and ClinVar annotations are shown with dark and light symbols; where conflicting, the MDSGene annotation was used. **B)** ROC AUC analysis showing the performance of MAVE data (green) and selected population free (cyan) and population tuned (grey) predictors in classifying benign vs pathogenic variants. **C)** Distribution of variant annotations in the combined MDSGene and ClinVar datasets (left) and in the acmgscaler-based reclassification of VUS (right). **D)** Pearson correlation matrix of MAVE pS65-Ub scores and selected VEPs.

To quantitatively assess predictive power, we performed receiver operating characteristic (ROC) analysis using our functional scores to distinguish pathogenic from benign variants. We achieved an area under the curve (ROC AUC) of 0.944 (95% CI: 0.905–0.978), demonstrating strong discrimination between pathogenic and benign variants (**Fig. 3B**). Using a threshold score of -0.58, the model exhibited 87.5% sensitivity with perfect specificity (100%) for the detection of pathogenic variants. We then applied acmgscaler [43] which translates functional scores into ACMG/AMP evidence strengths, to evaluate the potential impact of our data on variants of uncertain significance (VUS). Among 224 missense variants from MDSGene and ClinVar, 184 (81%) were initially classified as VUS. Our data provided evidence of pathogenicity or benignity for the majority of these variants, with 150 and 5 meeting BS3_moderate and BS3_supporting evidence levels, respectively; and 17, 0 and 1 meeting PS3_strong, PS3_moderate and PS3_supporting levels. Only 11 (6%) variants remained without additional support (**Fig. 3C**). These results demonstrate that our assay provides strong functional evidence capable of substantially reducing VUS burden when integrated within the ACMG framework.

To benchmark variant effect predictors (VEPs) against our experimental data and clinical annotations, we ranked 100 predictors by their correlation with our functional scores and by their ROC AUC for ClinVar/MDSGene pathogenicity classification. Among the predictors we tested, popEVE, iGEMME, CPT, and AlphaMissense correlated best with functional data (Pearson >= 0.51). CADD, a predictor frequently used for clinical interpretation, showed a lower correlation (Pearson = 0.32) (**Fig. 3D, Supplementary Fig. 3, 4**). Correlations improved slightly when using the absolute values of functional scores, consistent with the idea that deviations from wild-type activity in either direction may be relevant for pathogenicity. Despite moderate correlations with experiments, most VEPs correlated strongly with one another and many achieved excellent performance against ClinVar/MDSGene annotations, matching or even outperforming experiment-based classification (**Fig. 3B,D, Supplementary Fig. 3**). As expected, VEPs that correlated strongly with data also performed well in the clinical classification task, with GEMME only outperformed by SNPred and MetaRNN, two predictors whose performance might be overestimated due to the use of clinical data in training; such predictors trained on clinical data might not perform as well for previously unreported variants [44]. Despite overall high performance, VEPs tended to underestimate the pathogenicity of some variants, such as S65A **(Supplementary Fig. 3)**. Together, these results demonstrate that MAVEs and VEPs provide complementary information for the interpretation of pathogenic variants, and underscore the importance of re-evaluating commonly used VEPs as experimental functional data become available.

### Residual Parkin activity predicts age of onset in patients with biallelic mutations

Pathogenic homozygous or compound heterozygous variants in the *PRKN* gene are fully penetrant, with symptoms often manifesting before 40 years of age, referred to as early-onset PD, whereas heterozygous carriers typically remain unaffected but several studies suggest a slightly increased risk of late-onset PD compared with the general population. To assess how residual Parkin activity relates to clinical features, we calculated biallelic activity scores for 633 (367 excluding Het) patients from MDSGene. Wild-type alleles were assigned a score of 0; missense alleles were assigned the average functional score of the corresponding amino acid variant from our data; and structural or nonsense variants were assigned a score of -1, reflecting the average score of nonsense variants in our dataset. Each patient’s biallelic score was defined as the average of their two allele scores.

This analysis revealed a clear relationship between biallelic activity scores and the age at onset (**Fig. 4A**). Patients who developed symptoms around the age of 20 years typically carried two loss-of-function alleles with combined scores near -1; those with an onset between 40 and 50 years scored near -0.5 (e.g. one LOF allele and one milder missense variant). By contrast, patients with onset around 60-70 years – similar to the general population – had scores approaching 0, consistent with wild-type-like Parkin activity. A similar correlation was observed when analysing the average age at onset for each homozygous or compound heterozygous genotype individually, without binning patients by age (**Fig. 4B**). Including heterozygous cases reduced the strength of this correlation (**Fig. 4C**), likely due to misclassification: some of these individuals may in fact be undiagnosed compound heterozygotes carrying a second pathogenic PRKN variant - such as a structural rearrangement or deep intronic mutation - not routinely screened for in clinical genetics. These individuals might be erroneously treated as single heterozygotes, inflating their biallelic activity scores by assigning a functional wild-type allele instead of an undetected loss-of-function allele.

**Figure 4.**
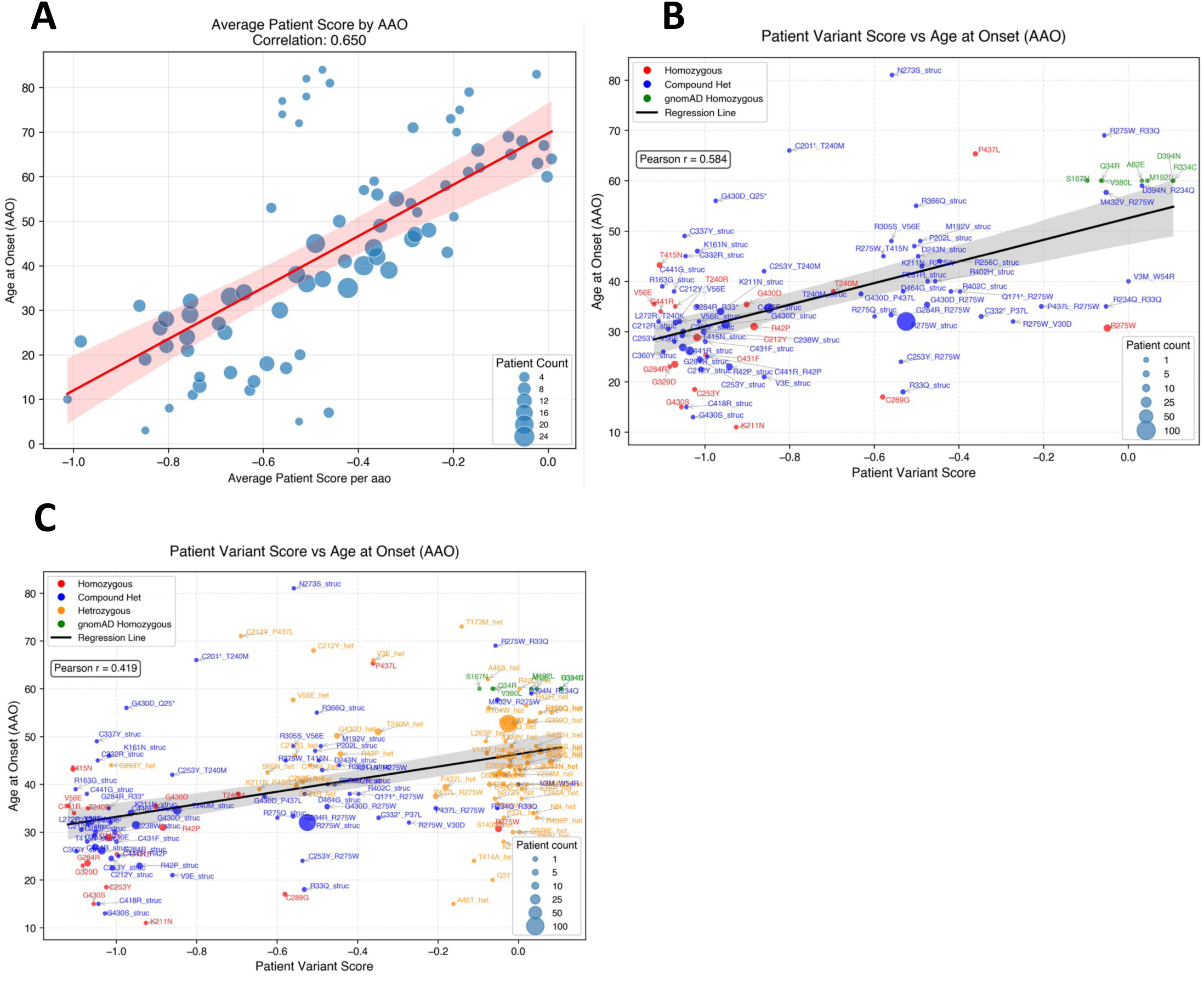
Prediction of PD age at onset (AAO) based on patient-level biallelic scores. **A)** Patients were grouped by age of onset, and mean biallelic scores were calculated for each group using the pS65-Ub score of each variant; the correlation between individual ages of onset and the corresponding group-averaged scores is shown. Point sizes show the number of patients in each group. **B)** Patients sharing the same genotype were grouped, and biallelic scores were calculated for each genotype; the correlation between average age of onset per genotype and the genotype’s biallelic score is shown. Point sizes show the number of patients in each group. Homozygous variants are shown in red, compound heterozygous variants in blue, and gnomAD homozygous variants (AF > 0.02; homozygous count ≥ 17) in green. **C)** Same as B), but including heterozygous variants (orange).

Notably, most individual *PRKN* variants had wild-type-like activity in our assay (**Fig. 2A, 2B**), and most variants in the general population are likewise expected to retain Parkin activity. Because these variants do not cause PD, their carriers are typically excluded from patient-oriented databases. Consistent with this, few patients in our dataset have biallelic variant scores near 0 (**Fig. 4B**), reflecting the rarity of wild-type-like genotypes in MDSGene. While this design supports accurate clinical annotation, it also leads to underrepresentation of functionally neutral variants, which may limit the use of such datasets for benchmarking high-throughput functional assays.

### Abundance profiling and comparison with VAMP-Seq

Clausen et al. [28] recently performed saturation mutagenesis of *PRKN* expressed in cells and measured variant effects on protein abundance using the fluorescence-based VAMP-Seq assay. Our functional MAVE scores show a moderate correlation with abundance (**Fig. 5**, **Supplementary Fig. 5**, Pearson = 0.46), potentially reflecting methodological differences: Clausen et al. expressed N-terminal GFP tagged Parkin vs our untagged Parkin; used a different cell line (HEK293 vs our HeLa); and did not include a mitochondrial damaging agent to trigger PINK1 activation and ensuant conformational changes and activation of Parkin via PINK1-mediated phosphorylation. To enable comparison, we used our pooled cell line to perform an abundance assay analogous to VAMP-Seq, using a fluorescent monoclonal antibody in place of EGFP as the readout. As in the pSer65-Ub assay, we included treatment with OA to induce mitochondrial damage.

**Figure 5.**
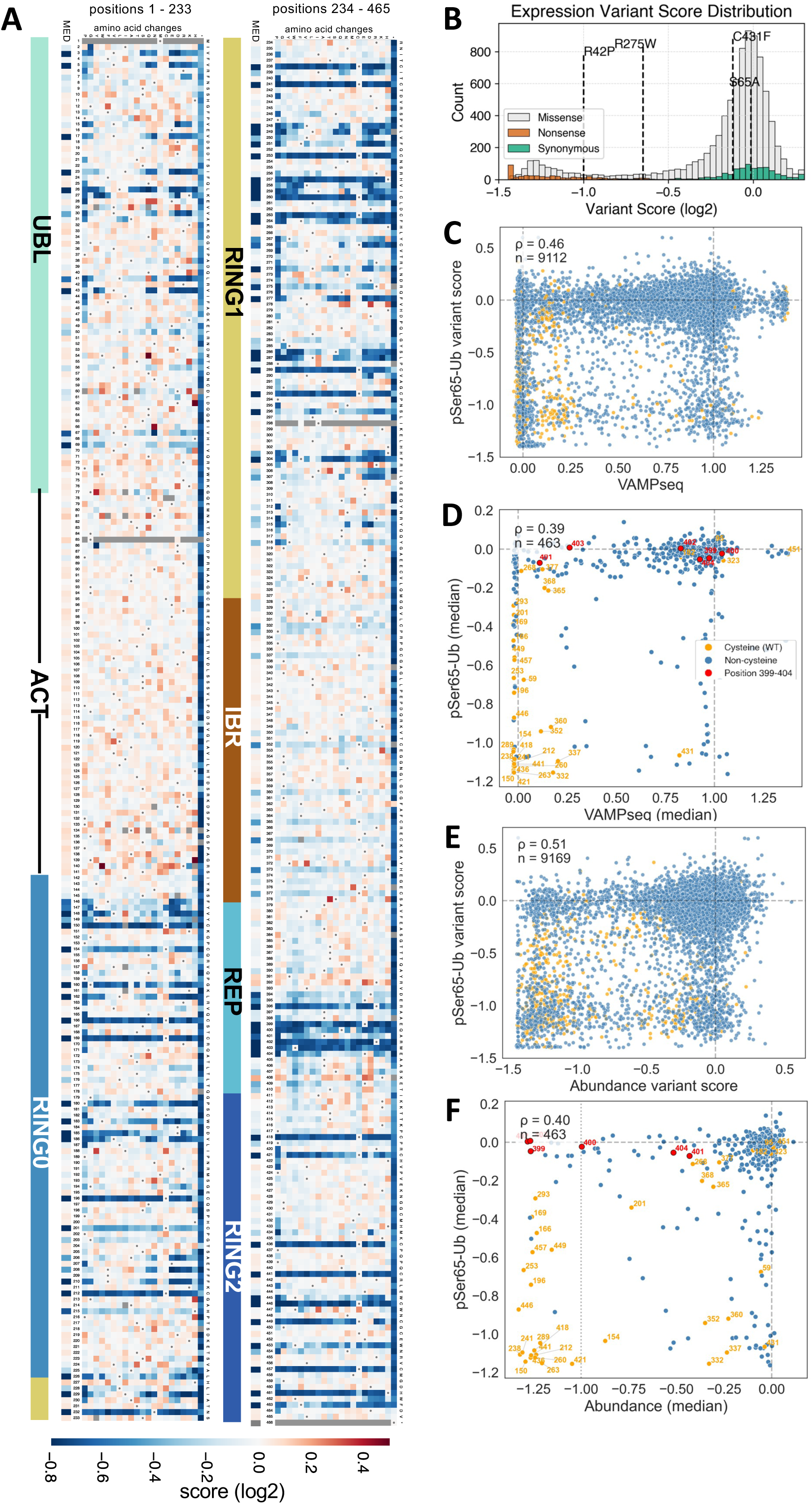
Comparison of abundance vs functional scores. **A)** Variant effect map for Parkin abundance. Colours as in Fig. 2A. **B)** Distribution of Parkin abundance scores for all missense (grey), synonymous (green) and nonsense (orange) variants. The median score of wild type barcodes is set as 0 and the median score of nonsense variants is -1. **C-F)** Correlation scatter plots between pS65-Ub functional scores, abundance scores and VAMP-Seq scores from (Clausen et al., 2024), as indicated. Orange points show cysteine residues, red points show the epitope of the monoclonal antibody used in our abundance assay (399-404) and blue points show other residues. C) and E) show variant scores, D) and F) show average variant scores per position.

We obtained abundance scores for 9,205 (99%) of missense and nonsense *PRKN* variants. As expected, synonymous variants scored similar to wild-type (average score = -0.02), stop codons ablated expression, and missense variants showed a bimodal distribution (**Fig. 5A,B**). Some variants showed divergent effects on function and expression: catalytic residues like C431 were essential for function but not for stability, while others (e.g. P180) were destabilizing but retained functional activity. Our abundance scores showed a higher correlation with VAMP-Seq (R=0.71) than with our functional scores (R=0.51), indicating consistent effects of variants on Parkin abundance across the two cellular models (**Fig. 5C,E, Supplementary Fig. 5**). The correlation decreased slightly when we compared the median score per position (**Fig. 5D,F**). The largest discrepancies with VAMP-Seq were seen at the N-terminus, where variants were more damaging in VAMP-Seq - possibly due to the N-terminal tag, which could have enhanced destabilisation and autoactivation of Parkin via removal of UBL-mediated autoinhibition [23]. Furthermore, residues 396-404 appeared more damaging in our assay, likely reflecting disruption of monoclonal antibody binding [26]. The three pathogenic variants with wild-type-like scores in our pSer65-Ub assay, R275W, R33Q and G328E, showed various degrees of destabilisation in our abundance assay and in VAMP-Seq. Interestingly, VAMP-Seq showed the highest correlation with VEPs, followed by our abundance assay and our pSer65-Ub assay (**Supplementary Fig. 3,6,7**). Nevertheless, protein abundance alone has limited power to distinguish benign from pathogenic variants (AUC ROC = 0.7 in our abundance assay and 0.78 in VAMP-Seq) or to reclassify VUS (**Supplementary Fig. 8**). Combining pSer65-Ub and abundance scores did not improve classification of benign vs pathogenic variants (**Supplementary Fig. 8**), but it increased the correlation of biallelic scores with the age at onset (**Supplementary Fig. 9**). Taken together, these results highlight that abundance- and activity-based assays capture complementary aspects of *PRKN* variant effects, suggesting that both may be required for clinical decision-making.

### Structural basis of Parkin variant effects

Over the past decade, multiple structures of Parkin and its activation intermediates have provided substantial insight into the molecular mechanism of Parkin activation [10, 11]. To elucidate the structural basis of the inactivating and activating variants identified in our screen, we mapped variants of interest (VOIs) onto the Parkin structure and interpreted their effects in the context of known regulatory mechanisms. This mapping revealed that inactivating variants clustered within key functional regions of Parkin, where they are likely to impair activity by destabilising the folded protein structure, perturbing catalytic residues, or disrupting essential interdomain interfaces required for activation (**Fig. 6 & Supplementary Fig. 10A-G**). Conversely, variants enriched in regulatory regions that relieve autoinhibition were frequently activating (**Fig. 6 & Supplementary Fig. 10F**).

**Figure 6.**
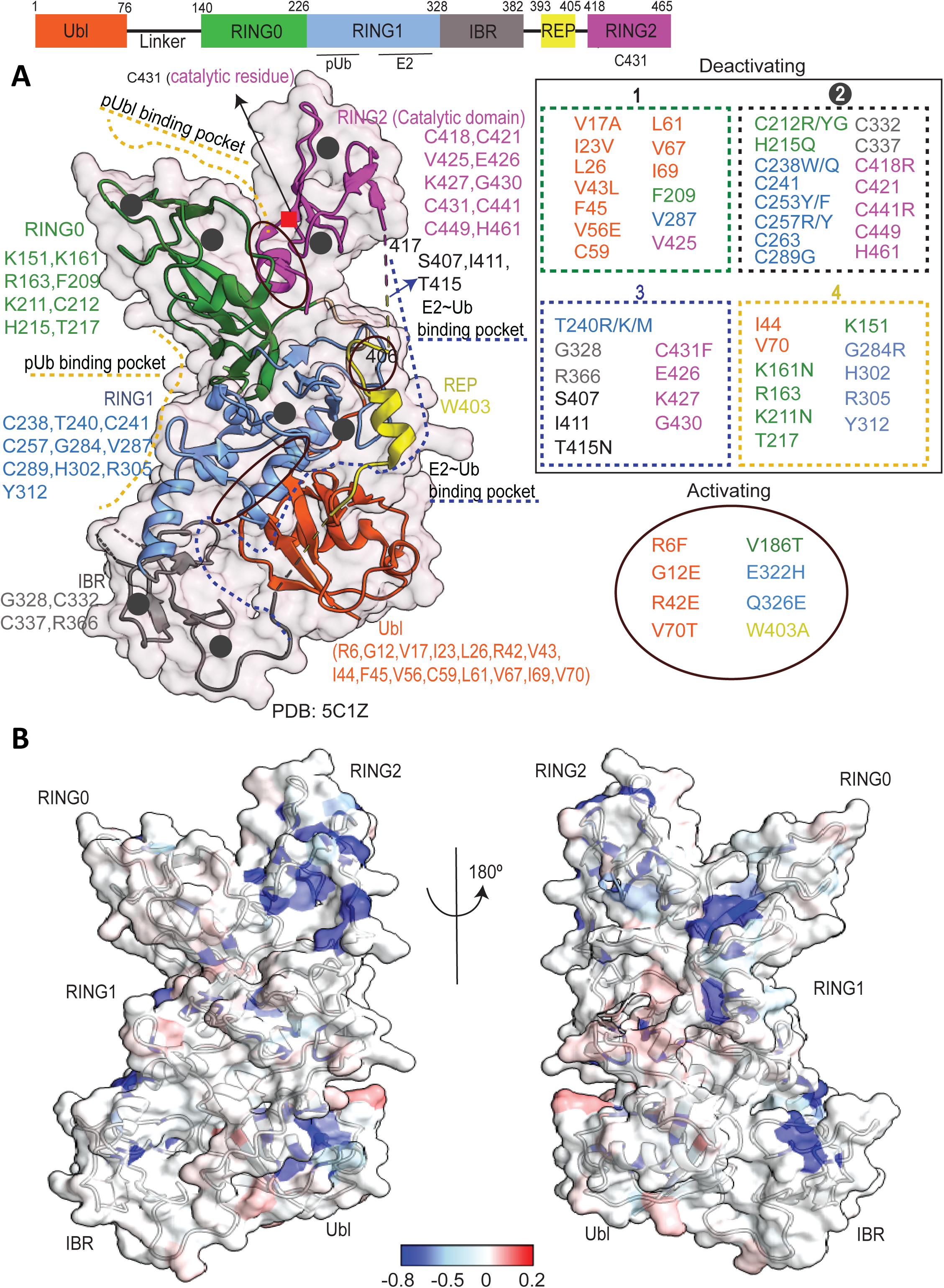
Structural interpretation of Parkin variants. **A) (Left)** Schematic representation of Parkin domain organization (upper panel) and the crystal structure of autoinhibited Parkin (lower panel), with key functional elements annotated. **(Right)** Selected inactivating and activating variants grouped according to their proposed mechanisms of action. Residues are colored by domain, with residues from disordered regions shown in black. **B)** Parkin structure coloured by the median functional score of mutations per position, using a colour scale as in Fig. 2

We first examined positions at which the majority of amino acid substitutions were inactivating, revealing four major mechanistic classes (**Fig. 6A, Supplementary Fig. 10A-G**): (1) variants that destabilise the protein by disrupting hydrophobic core packing; (2) variants that compromise Zn²⁺ coordination and thereby destabilise Parkin; (3) variants that impair catalytic activity by perturbing E2∼Ub binding or transthiolation; and (4) variants that impair activation by disrupting regulatory interfaces, including contacts with PINK1, phospho-ubiquitin, or phospho-UBL.

Class-1 inactivating variants were distributed across all Parkin domains, with prominent clusters in the UBL, RING0, RING1, and RING2 domains (**Fig. 2A**, **Fig. 6A, Supplementary Fig. 10A**). Disruption of hydrophobic cores, particularly within the UBL and RING0 (**Supplementary Fig. 10A**), generally reduced both Parkin activity and protein abundance (**Fig. 2A**, **Fig. 5A**), consistent with their significance in protein stability. However, certain core mutations near the UBL phosphorylation site (e.g. C59, L61) abolished activity without affecting abundance, likely by interfering with PINK1 binding and subsequent Parkin phosphorylation (**Supplementary Fig. 10G**).

Class-2 variants affecting Zn²⁺-coordinating residues clustered within the RING0, RING1, IBR, and RING2 domains (**Fig. 6A, Supplementary Fig. 10B**). While most of these residues were essential for both activity and stability, a subset, particularly residues in the second Zn²⁺-binding motif of the IBR domain (C365, C368, H373, C377), tolerated mutation in both functional and abundance assays. This observation is consistent with the relatively dynamic or partially disordered nature of the IBR domain observed in Parkin structure showing a higher B-factor (**Supplementary Fig. 10H**).

Class-3 variants that impaired catalytic activity, including substitutions at the E2∼Ub interaction interface (e.g. S407, I411, T415) [37] or at the active-site cysteine (C431) (**Fig. 6A, Supplementary Fig. 10C**), typically retained near-wild-type protein abundance despite exhibiting severe loss of function. This indicates that these variants primarily disrupt enzymatic activity rather than protein stability.

Class-4 variants affecting various regulatory interfaces constituted the most heterogeneous group. These alterations disrupted intermolecular interactions with PINK1 (e.g. I44) or phospho-ubiquitin (e.g. R163), as well as intramolecular regulatory interfaces, including pUBL–RING0 (e.g. K211), and RING0-ACT (e.g. L102) interfaces (**Fig. 6A, Supplementary Fig. 10D-E**). Some residues participate in multiple regulatory interactions; for example, I44 mediates both PINK1–UBL binding (**Supplementary Fig. 10G**) and UBL–RING1 autoinhibition (**Supplementary Fig. 10F**). Accordingly, the I44A variant disrupts Parkin phosphorylation by PINK1 while simultaneously relieving autoinhibition [23, 45]. In our dataset, most I44 substitutions were loss-of-function, with the exception of I44F, which was activating (**Fig. 2A; Supplementary Table 2**). In contrast, alterations at K211 abolished Parkin activation by preventing pUBL binding to RING0 (**Supplementary Fig. 10E**); correspondingly, all K211 substitutions except K211R were inactivating in our assay. Overall, mutations at regulatory interfaces frequently exhibited complex behaviour, with different substitutions at the same position producing either strong loss- or gain-of-function effects.

We next analysed positions where most substitutions increased Parkin activity. These sites were common in the ACT, RING0, RING1, and REP domains, where 28–45% of residues showed net activating behaviour (functional score > 0) (**Fig. 2A**). The strongest activating effects were concentrated in the UBL domain: eight of the top eleven residues ranked by median functional score were located there, predominantly in charged residues (R6, E16, R42, K48, R51, R72, R75) (**Fig 5A, Supplementary Fig. 10F**). Several of these activating hotspots have been reported previously. In particular, R6, R42, K48, V70, and R72 are key residues at the UBL autoinhibitory interface, and their substitution promotes Parkin activation [23, 46]. Notably, whereas prior studies were performed in the absence of Parkin phosphorylation, our experiments, conducted in a cell-based system with phosphorylated Parkin, revealed contrasting effects for certain variants, such as R42P (loss-of-function) and V70T (gain-of-function). We believe that these differences arise from the unique positioning of R42 and V70 at the interface between pUBL and RING0 (**Supplementary Fig. 10E**), where substitutions may differentially modulate this interaction and thus Parkin activation. Moreover, both residues also contribute to the PINK1–UBL binding interface [47] (**Supplementary Fig. 10G**), suggesting that they may additionally influence Parkin phosphorylation efficiency, further contributing to their distinct functional outcomes.

## Discussion

Here we systematically measured the effects of nearly all possible missense and nonsense variants in *PRKN* using a pooled cellular assay of phosphorylated ubiquitin (pSer65-Ub). The resulting map identifies loss- and gain-of-function variants across the protein; resolves the functional impact of hundreds of VUS; and reveals a quantitative relationship between residual Parkin activity and age at onset for Parkinson’s disease (PD).

Historically, a wide range of assays have been used to study Parkin function and interpret the impact of disease-associated variants. These have included in *vitro assays* of recombinant Parkin that enable direct measurement of catalytic activity based on autoubiquitylation or substrate-based ubiquitylation [23, 24]; human cell based assays of Parkin ubiquitylation, pSer65-Ub, and mitophagy [25, 26, 42] as well as in vivo models in Drosophila and mice [42, 48, 49]. HeLa cells have been extensively used in over-expression studies of Parkin function in cells since they do not express endogenous Parkin thereby allowing clear-cut dissection of the effect of mutant Parkin compared to the wild-type [50, 51]. Furthermore, mechanistic understanding of Parkin function in HeLa cells has been validated in neuronal systems [42, 52]. Each assay captures different aspects of Parkin activity e.g. the pathogenic R42P variant [53] exhibited no effect via *in vitro* ubiquitylation assays [23, 40], but was shown to disrupt folding [54], and reduce Parkin function in cell-based assays [25], the latter consistent with our study.

Furthermore, in another study, Y143D, Y143E, A401K and F463K variants strongly activated Parkin *in vitro* but did not increase mitophagy in cells [26], and consistent with this we found that these variants also had minor effects on Parkin activation in cells. This illustrates how different assays can report distinct effects depending on whether they measure protein stability, biochemical activity, or more downstream cellular phenotypes.

In parallel to our study, the Hartmann-Petersen group performed a comprehensive variant effect screen of Parkin using a mitophagy-based assay, generating functional scores for >9,200 variants [55]. Despite differences in cellular models, functional readouts, and library designs, the results of the two screens were remarkably similar (R=0.77) and both screens performed well at distinguishing benign from pathogenic Parkin variants (ROC AUC 0.92 and 0.94 for the mitophagy and pS65-Ub assays, respectively). Classification of benign vs pathogenic variants was markedly more accurate in the two functional assays than in abundance-based assays performed by the same groups (ROC AUC 0.78 and 0.7 for VAMP-Seq [28] and our antibody-based assay). This suggests that MAVEs with bespoke functional readouts can not only provide mechanistic information about gene function, but also perform well in pathogenicity prediction.

While MAVEs often identify only a few activating variants, we observed an unusually high prevalence in Parkin: nearly 40% of residues in the ACT, RING0, RING1, and REP domains showed more activating than inactivating substitutions; 2.5% of all substitutions and 4.5% of substitutions in the UBL domain were significantly activating (FDR<0.05). In some previous MAVE studies of other genes, gain-of-function effects were linked to increased substrate affinity [56, 57] or elevated protein expression [30]. The high numbers of activating mutations in Parkin, identified here, could be due to its structural regulation via autoinhibition: most activating variants were in autoinhibitory interfaces, and many different substitutions at these interfaces might be expected to relieve autoinhibition. Similar mechanisms explained activating mutations in the human glucokinase GCK [58] and in the DNA methyltransferase DNMT3A [59]. Detection of activating variants could also have been facilitated by our experimental design, which captured pS65-Ub 45 minutes after mitochondrial damage. By comparison, Sigmarsdóttir et al. [55] used GFP-tagged Parkin and measured mitophagy 4 hours after OA treatment, potentially masking early kinetic effects of mutations.

Previous targeted structure-function studies found activating variants mainly clustering around the REP:RING1 (e.g. W403A) or RING0:RING2 (e.g. F146A) interfaces that were also active in cells at promoting mitophagy after 4 hours of mitochondrial damage [26]. Interestingly, we did not observe strong activation of W403A or F146A variants suggesting that these variants may have more specific downstream effects on mitophagy not captured by our assay conditions. Activating variants, though mechanistically distinct from loss-of-function variants, can still be pathogenic and may demand entirely different therapeutic approaches. Because current VEPs poorly predict gain-of-function effects, MAVEs remain essential for their discovery and interpretation.

R275W represents one of the most common pathogenic variants of Parkin; it is found more often as a compound heterozygote and more rarely as homozygous variant in PD cases [22, 33, 34]. It is also frequently found in control databases as a heterozygous variant. Previous in vitro studies of recombinant Parkin have shown that the R275W variant reduces Parkin activity [40, 60]. Furthermore, the recent VAMP-Seq study of GFP-Parkin found that R275W led to a drastic reduction of Parkin abundance [28]. We found that R275 variants had only mild effects on Parkin activity, and moreover we observed that the R275W variant led to a moderate decrease in Parkin abundance. It is therefore likely that under our assay conditions of stable Parkin over-expression, the effect of the decreased expression of Parkin on activity is masked by feed-forward amplification. The effect of R275W on Parkin stability that we observe is consistent with biochemical analysis of primary cells derived from PD patients harbouring R275W compound heterozygous mutations, in which it was found that the R275W mutation led to lower but not complete loss of total endogenous Parkin expression and activity [61].

The role of heterozygous PRKN mutations in influencing disease risk and manifestation is a matter of ongoing debate. Large-scale genetic screens—such as a meta-analysis of 19,574 cases and 468,488 controls [62] and a population-based recall-by-genotype study from the CHRIS cohort showing high carrier frequencies but low penetrance of PD risk markers [63]—found no or only marginally increased PD risk in monoallelic carriers, a conclusion reinforced by UK Biobank and AMP-PD data [64]. However, these studies generally lack deep clinical phenotyping of controls, leaving subtle motor and non-motor signs undetected. Notably, monoallelic PRKN pathogenic variant carriers have a younger age at onset than non-carriers and heterozygous copy number variants may confer a higher risk than single nucleotide variants [62], indicating that both mutation type and variant severity modulate penetrance even in the heterozygous state. In addition, carefully designed studies in smaller cohorts—typically examining the offspring of biallelic mutation carriers who are unlikely to harbor a second, overlooked variant—have consistently revealed preclinical abnormalities, including reduced striatal ^18^F-dopa uptake on PET, substantia nigra hyperechogenicity, and electrophysiological changes [65]. Moreover, sensor-based gait and posturography assessments under challenging motor conditions, performed blinded to the genetic status, discriminated clinically unaffected heterozygous carriers from mutation-free controls with up to 87% accuracy, even when standard neurological examination showed no differences [66]. These findings suggest that heterozygous PRKN variants produce a subclinical vulnerability that, depending on variant severity and additional genetic or environmental hits, may push carriers across the threshold of disease manifestation. The availability of functional data for thousands of Parkin variants will increase the power to detect subtle changes in PD risk.

Our functional map of Parkin variants offers immediate clinical utility for variant interpretation, enabling classification of the vast majority of VUS and improving diagnostic certainty for patients undergoing genetic testing. Beyond diagnosis, the quantitative relationship between genotype scores and age at PD onset establishes a framework for individualized prognostication. This genotype-phenotype correlation is exemplified by recent work [67], demonstrating that patients carrying homozygous PRKN Exon 2 deletions experience disease onset approximately 13 years later than other PARK-PRKN patients, attributed to residual ParkinΔ1–79 activity that can be pharmacologically enhanced by the molecular glue BIO-2007817. Critically, this same modulator inhibited wild-type full-length Parkin, underscoring the need for genotype-based patient stratification in therapeutic development. While our data currently predict age at onset, future integration with longitudinal cohorts may enable prediction of additional clinical outcomes including motor and cognitive progression, non-motor symptom burden, and response to dopaminergic or disease-modifying therapies—outcomes that await systematic collection of deep clinical phenotyping data linked to functional variant scores.

## Methods

### General experimental reagents

Enzymes were purchased from New England Biolabs unless otherwise specified. Oligonucleotides were synthesized by IDT and are listed in **Supplementary Table 3**. HeLa cells carrying the G542A_pLenti-Tet-coBxb1-2A-BFP_IRES_iCasp9-2A-Blast_rtTA3 landing pad were cultured in Dulbecco’s Modified Eagle’s Medium (DMEM) (Thermo Fisher Scientific, 41965039) supplemented with 10% fetal bovine serum and Penicillin–Streptomycin (50 I.U./mL). Cells were passaged by trypsinization using 0.25% trypsin–EDTA (Millipore-Sigma) and tested negative for Mycoplasma before use. All antibodies used are detailed in **Supplementary Table 4**.

### HeLa BxB1 cell treatment and preparation for pS65 Ubiquitin and total Parkin flow cytometry

HeLa Bxb1 cells were grown and maintained in DMEM supplemented with 10% fetal calf serum, 2 mM L-glutamine, penicillin (100 U/mL), streptomycin (100 µg/mL), doxycycline (3 µg/mL; Sigma), and puromycin (1 µg/mL). All cells were maintained at 37 °C in a humidified atmosphere containing 5% CO₂. Cells were detached with trypsin, counted, and resuspended at a density of 2.5×10⁶ cells/mL in culture medium without puromycin. To stimulate Parkin activity, cells were incubated in suspension with 1 µM oligomycin A (Sigma, 75351) and 10 µM antimycin A (Sigma, A86740) for the indicated times at 37°C in 5% CO₂. For oligomycin and antimycin time-course experiments, treatments were applied at 10-min intervals over an 80-min period. An incubation time of 45 min was used for sorting HeLa BxB1 cells expressing the Parkin library. Following incubation, cells were washed twice with ice-cold PBS containing 0.5 mM EDTA. Cells were incubated with ice-cold Aqua Live/Dead fixable dead cell stain (Thermo Fisher Scientific) for 10 min, followed by two additional washes with ice-cold PBS containing 0.5 mM EDTA.

For the pS65 Ubiquitin assay, cells were fixed with 2% PFA for 10 min and washed twice with PBS before permeabilisation. Permeabilisation was performed by incubating cells in 90% methanol for 30 min at −20 °C. Cells were washed twice with PBS containing 1% FBS and resuspended in PBS containing 1% BSA to a final density of 5×10⁶ cells/mL. Cells were incubated with the rabbit monoclonal phospho-ubiquitin (Ser65) antibody (Cell Signalling Technology, #62803) at a final concentration of 1.25 ng/µL for 1 hr at room temperature. Cells were then washed twice with PBS before incubation with the goat anti-rabbit-Alexa Fluro 647 antibody (Thermo Fisher Scientific, #A21244) at a final concentration of 4 μg/mL for 1 hour at room temperature. Following incubation, the cells were washed a final 2x before resuspension in PBS containing 1% FBS.

For the total Parkin assay, cells were fixed with 2% PFA for 20 min and washed twice with PBS before permeabilisation. Permeabilisation was performed by incubating cells at room temperature in 1× permeabilisation buffer diluted in PBS (eBioscience, Thermo Fisher Scientific) for 30 min. Cells were washed twice with PBS containing 1% FBS and resuspended in PBS to a final density of 5×10⁶ cells/mL. Cells were incubated with the mouse monoclonal total Parkin (PE-conjugated) antibody (Santa Cruz Biotechnology, sc-32282-PE) at a final concentration of 5 µg/mL for 1 hour at room temperature. Following incubation, the cells were washed a final two times before resuspension in PBS containing 1% FBS.

Within 24 hours of preparation, samples were run on the NovoCyte (ACEA Biosciences) for optimisation, and subsequently single-cell sorted using the FACS Aria II.

### HeLa BxB1 cell treatment and lysis

HeLa cells were cultured in DMEM supplemented with 10% fetal calf serum, 2 mM L-glutamine, penicillin (100 U/mL), streptomycin (100 µg/mL), and doxycycline (3 µg/mL; Sigma). To stimulate Parkin activity, cells were incubated for 45 minutes with oligomycin A (Sigma, 75351) and 10 µM antimycin A (Sigma, A86740) at 37°C with 5% CO_2_. Following incubation, cells were rinsed with ice-cold PBS, and lysed in ice-cold lysis buffer (50 mM Tris-HCl (pH 7.5), 250 mM sucrose, 1 mM EDTA, 1 mM EGTA, 1mM sodium orthovanadate, 10 mM sodium β-glycerophosphate, 50 mM sodium fluoride, 5 mM sodium pyrophosphate, 1% (v/v) Triton-X 100, supplemented with CompleteTM EDTA-free protease inhibitor cocktail (Roche, 11836170001), phosSTOPTM (Roche, 4906845001), and 200 mM chloroacetamide (Sigma-Aldrich). Cell lysates were snap frozen and maintained at −80 °C.

### Immunoblotting

Protein lysates were clarified by centrifugation at 17,000×g for 20 min at 4 °C. Protein concentration was determined from the resulting supernatant using the Pierce BCA Protein Assay Kit (Thermo Fisher Scientific). Cell lysates were mixed with the NuPAGE LDS Sample Buffer (Life Technologies) supplemented with 5% (v/v) 2-mercaptoethanol and heated to 90 °C for 3 min. 5-10 μg of protein was loaded onto NuPAGE 4–12% Bis-Tris Midi Gels (Thermo Fisher Scientific, WG1403BX10). Samples were electrophoresed in NuPAGE MOPs SDS running buffer (Thermo Fisher Scientific), following the manufacturer’s instructions. Protein transfer was performed on ice onto nitrocellulose membranes (GE Healthcare, Amersham Protran Supported 0.45 mm NC) at 100 V for 90 min in transfer buffer (48 mM Tris–HCl and 39 mM glycine supplemented with 20% methanol). Membranes were then incubated with 5% (w/v) skim milk dissolved in TBS-T (20 mM Tris–HCl, pH 7.5, 150 mM NaCl, and 0.1% (v/v) Tween 20) at room temperature for 45 minutes. Primary antibodies were prepared in 5% BSA (Sigma) dissolved in TBS-T. Membranes were washed once before incubation overnight at 4 °C with primary antibodies. Following incubation, membranes were washed with TBS-T for 5 minutes 3x. Secondary antibodies were prepared in 5% skim milk dissolved in TBS-T. Membranes were incubated for 45 minutes at room temperature with secondary antibodies. Membranes were washed with TBS-T for 10 minutes (3x). Image acquisition was performed with near-infrared fluorescent detection using the Odyssey CLx imaging system and quantified using the Image Studio software.

Anti-PARKIN (PARK2) sheep polyclonal antibody, purified by MRC PPU Reagents and Services at the University of Dundee (S966C), was used at a 1:2000 dilution. Anti-Parkin phospho-Ser65 rabbit monoclonal antibody, generated by Epitomics/Abcam in collaboration with the Michael J. Fox Foundation for Research, was used at a 1:2000 dilution. Anti-Ubiquitin phospho-Ser65 (E2J6T) rabbit monoclonal (Cell Signalling Technology, 628002S) and anti-PARKIN (PRK8) mouse monoclonal antibody (Santa Cruz Biotechnology, sc-32282) were used at 1:2000 and 1:1000, respectively. Anti-GAPDH (6C5) mouse monoclonal antibody (Santa Cruz Biotechnology, sc-32233) was used at 1:10,000. The secondary antibodies, donkey anti-goat (LI-COR, 926-68074), donkey anti-rabbit (LI-COR, 926-32213), and donkey anti-mouse (LI-COR, 926-32212) were used at a 1:20,000 dilution.

### Plasmid construction

The human PRKN coding sequence (UniProt O60260-1) was subcloned from pcDNA5D-FRT/TO/PRKN[WT] (MRC PPU Reagents and Services University of Dundee, DU23307). PRKN coding sequence from pcDNA5D-FRT/TO/PRKN[WT] was amplified using VA01/VA02 primers followed by restriction digestion by KasI and EcoRI. The digested product was then purified and cloned into a Gateway entry vector pAINt (Çubuk et al., 2025). Vector plasmid is also digested by the same enzymes and dephosphorylated by Shrimp Alkaline Phosphatase (rSAP). attB_PRKN_IRES_PuroR expression plasmids were generated via Gateway LR recombination (Thermo Fisher Scientific, 11791020) between the pAINt-PRKN_entry plasmids and attB_ccdB_IRES_PuroR (pMV2) destination plasmid. Correct plasmid sequences were confirmed by whole-plasmid sequencing.

### Generating barcoded destination vector

The pMV2 destination plasmid (16μg) was digested by adding 80μL of CutSmart Buffer (10x), 8uL of SacI and 8μL of XhoI, and the final volume was brought up to 800μL with nuclease free water. This was gently mixed and then divided into eight tubes and incubated overnight at 37℃, and dephosphorylated by adding 3μL of rSAP into each tube and incubation at 37℃ for 30 minutes. The reaction was cleaned by QiaQuick PCR Cleanup Kit (Qiagen, 28106) by running it through a single column to limit DNA loss. pMV2_BC barcoding oligos flanked by SacI and XhoI sites were synthesized and PAGE-purified before pMV2_BC_R primer annealing and extension using Phusion HiFi PCR master mix (M0531S). 16µL of the barcoding oligo (10µM) was combined with 40µL of pMV2_BC_R oligo (10µM), 400µL of 2× Phusion Master Mix and 344µL nuclease-free water. This was gently mixed and then divided into 50µL aliquots in PCR tubes and run with the following PCR conditions in the thermocycler: 98℃ for 2 minutes, 60℃ for 30 seconds and 72℃ for 60 seconds. The reactions were then cleaned with the Oligo Clean & Concentrator kit (Zymo Research, D4061), quantified and 200ng was digested with 1.25µL SacI and 1.25µL XhoI, 10µL CutSmart Buffer (10×), with the final volume being brought up to 100µL with nuclease-free water. This digestion was incubated at 37℃ overnight and cleaned again with the Oligo Clean & Concentrator kit. The ligation was performed as a 1:1 ratio mixing 6µg of pMV2 plasmid with 92.4ng of the digested barcode with adding 92µL of T4 Ligase buffer, 34.5µL of T4 DNA Ligase, and the final volume was brought up to 920µL with nuclease free water. The reaction was gently pipetted up and down to mix and was divided into 20µL aliquots for incubation at 4℃ for 72 hours before combining and transferring into two low bind tubes (Eppendorf). We used six 0.2cm gap electroporation cuvettes (BioRad) to transform the barcode library into the Electro-Competent ccdB Survival cells. Two tubes of 500µL of the competent cells were thawed on ice for 10 minutes and transferred to the reaction tubes, gently mixed with the pipette, avoiding making bubbles, and then 100µL at a time was transferred to the electroporation cuvettes and electroporated (Bio-Rad GenePulser at 2.5 kV, 200 Ω, and 25 μF), and transferred to 100mL of prewarmed SOC in a 500mL flask. This process was repeated until all of the cells had been electroporated and added to the same flask of SOC. The SOC flask was then rotated at 37℃ for 1 hour. Next, 10µL of the mixture along with 1/100 and 1/10000 dilutions plated on LB carbenicillin plates. The rest of the 100mL of barcode transformation was moved to 1L of 100µg/mL carbenicillin LB and incubated at 30℃ 200rpm for about 19 hours, or until an OD600 of about 0.5 was reached. Cells were then spun down at 4000×g for 15 minutes, resuspended in a small amount of LB and aliquoted into 1.5mL tubes and stored as pellets after centrifugation -20℃. We estimate having around 20 million unique barcodes by counting colonies on the dilution plates. Several colonies from plates were sequenced to ensure proper ligation.

### Making Electro-Competent ccdB Survival Cells

An antibiotic-free LB Agar plate was streaked with One Shot^TM^ ccdB Survival^TM^ T1^R^ Competent Cells (Invitrogen) and grown overnight at 37℃. Super Optimal Broth (SOB) was prepared: 20 g tryptone, 5 g yeast extract, 0.5 g sodium chloride, 0.186 g potassium chloride, made to 1 L with MilliQ Water, and autoclaved the same day. After autoclaving, 5mL of 2M MgCl_2_ (w/v) was added. The following day, a single colony was picked from the overnight plate and inoculated into 5mL of SOB and grown overnight at 37℃ 200rpm. The following day, the entire 5mL overnight culture was added to 1L of SOB in a 5L flask. This was incubated at 18℃ 200rpm and grown for about 36 hours, or until the OD600 reached 0.4. The 1L culture was then moved to an ice bucket and put on a rocker in the cold room for half an hour to fully cool. The culture was then divided between four, 250mL, sterile bottles and spun down at 4000×g for 15 minutes at 4℃. The supernatant was removed and each cell pellet was gently resuspended in 200mL of ice cold water. These were spun down at 4000×g for 10 minutes at 4℃. Another 200mL ice cold water wash and spin down was performed. The supernatant was removed and each pellet was resuspended in 100mL of ice cold 10% glycerol (v/v) and combined into two bottles. These were spun down one more time at 4000×g for 15 minutes at 4℃. The supernatant was carefully removed, ensuring that the glycerol-loosened pellet was not disturbed. The final pellets were lifted and combined with about 3.5mL of 10% glycerol. With the volume of the pellet, this resulted in about 15mL of competent cells. The pellet was divided into 500µL aliquots and snap frozen in liquid nitrogen, before being stored in the -80℃ freezer.

### Oligonucleotide pool design and preparation

We used SUNi primer design method [68] to design a combination of NNS and NNK degenerate oligonucleotides for introducing single amino acid mutations at each of the 464 codons of the *PRKN* coding sequence (excluding the start codon) (**Supplementary Table 3**). The NNS/NNK codon encodes all 20 amino acids and one stop codon. To prevent unequal representation of variants, the oligonucleotides were organized into 10 pools, each covering 42–47 sequential codons, and mutagenesis reactions were performed separately for each pool. Each oligonucleotide mixture was phosphorylated by combining 20 µL of 10 µM oligonucleotides with 2.4 µL T4 Polynucleotide Kinase buffer, 1 µL of 10 mM ATP, and 1 µL T4 Polynucleotide Kinase (10 U/µL). The reaction was incubated at 37 °C for 60 minutes. Phosphorylated oligos were stored at −20°C and diluted 1:500 in nuclease-free water on the day of mutagenesis. The universal secondary primer (VA03) was phosphorylated in the same manner using 7 µL of a 100 µM stock in a 30 µL reaction, and diluted 1:20 prior to use.

### Variant library generation by saturation mutagenesis

To construct the PRKN variant library, we used the nicking mutagenesis protocol [31]. The pAINt-PRKN plasmid used as template was freshly purified from bacterial pellets using the QIAprep Spin Miniprep Kit (Qiagen, 27106). The resulting mutagenesis product (pAINt-PRKN library) was eluted in 11 µL of nuclease-free water. Two transformations were performed, each using 5.5 µL of the mutagenesis product and 50 µL of NEB^Ⓡ^ 5-alpha competent *E. coli* (C2987H). Serial dilutions (1:10 and 1:100) of the transformation mixture were plated to estimate transformation efficiency and estimate number of variants. The remaining transformation mixture was inoculated into 100 mL LB medium with 50 mg/mL kanamycin and cultured overnight at 37 °C, 250 rpm. Cells were collected by centrifugation, washed with sterile water, re-pelleted, and aliquots were stored as pellets at −20 °C. Transformation efficiency inferred from dilution plates indicated an estimated 7.44 × 10⁵ colony-forming units (CFU) per pool.

### Generating barcoded PRKN variant library by Gateway cloning

Barcoded PRKN variant library was generated using Gateway LR recombination between the pAINt-PRKN plasmid library and barcoded attB_ccdB_IRES_PuroR (pMV2) destination plasmids. Separate LR reactions were performed for each of the 10 pools. For each reaction, 150 ng of entry plasmid and 150 ng of destination plasmid were combined with 2 µL of Invitrogen™ Gateway™ LR Clonase™ II enzyme mix (Thermo Fisher Scientific, 11791020) and TE buffer (pH 8) to a final volume of 10 µL. Reactions were incubated overnight at 25 °C, followed by addition of 1 µL Proteinase K (2 µg/µL) and incubation at 37 °C for 10 minutes to terminate the reaction. For each pool, two transformations were performed, each with 5 µL of the LR reaction product into 50 µL of NEB^Ⓡ^ 5-alpha competent *E. coli*. A 50 µL aliquot of the transformation mixture was plated on LB–agar supplemented with 100 µg/mL ampicillin to estimate transformation efficiency and number of barcoded variants. The remaining transformation mixture was inoculated into 100 mL LB medium containing 100 µg/mL ampicillin and cultured overnight at 37 °C, 250 rpm. Transformant counts from plates indicated that each pool contained approximately 20,000–24,000 CFU. Bacterial cultures were harvested and stored as pellets at −20 °C. Plasmid DNA was prepared from each pool using the Qiagen Miniprep Kit, and 50 µg of purified plasmid from each pool was combined to generate the final barcoded, mutagenized PRKN library.

### PacBio sequencing of barcoded PRKN variant library

PacBio long-read sequencing was performed to link each barcode to its corresponding PRKN variant. Sequencing templates were generated from the purified plasmid library by double digestion with AvrII and EcoRI-HF. The digest produced a 1.9 kb fragment containing the barcode and PRKN sequence, along with 3.7 kb, 261 bp, and 75 bp fragments corresponding to the plasmid backbone. The digestion products were purified using the QIAquick PCR Purification Kit and submitted to Edinburgh Genomics (University of Edinburgh) for size selection using BluePippin (Sage Science), followed by library preparation and sequencing on a single SMRT Cell using the PacBio Revio platform. We processed HiFi reads (raw sequencing files available in GEO; accession number GSE000000) using alignparse (Crawford and Bloom, 2019) version 0.6.3, which applies minimap (Li, 2018) version 2.24 for long read alignment to identify and call mutations in the PRKN sequence and to phase them with barcodes. 4,096,459 of 4,708,155 HiFi reads (87%) mapped to the target (available in GEO; accession number GSE000000), and the output was used to generate a codon-variant lookup table. We first retained mapped HiFi reads with sequencing accuracy reported by the PacBio ccs program in both PRKN sequence and barcode to be at least 99.99%. 68% of all reads passed these filters. As PacBio sequencing produced on average 30 reads per amplicon (**Supplementary Table 1**), we next calculated the empirical accuracy of retained reads (Starr et al., 2020) to assess the reliability of barcode-variant phasing, i.e., how often the reads linked with a specific barcode are identical. We observed 99.28% empirical accuracy for each HiFi read, excluding reads with indels. Since 76.14% of the barcodes were associated with two or more HiFi reads we generated a cumulative consensus of HiFi reads within these barcodes, so that consensus accuracy would exceed calculated empirical accuracy for individual reads. Barcodes associated with insertions, deletions or more than one variant were also excluded. Overall 609,187 barcodes were retained, of which 438,159 associated with 9,236 missense variants, 16,488 barcodes with 456 nonsense variants, 24,235 and 146,793 barcodes with synonymous variants and wild type sequence, respectively (**Supplementary Table 1**). The code used to generate the final barcode-variant table is available on Github () and includes additional plots illustrating library structure.

### HeLa landing pad cell lines generation

A previously reported doxycycline-inducible BxB1 recombinase landing pad construct (Tet-coBxb1-2A-BFP_IRES-iCasp9-2A-Blast_rtTA3; Addgene #171588) was delivered into HeLa cells via lentiviral transduction using the Lenti-X Packaging Single Shot (VSV-G) system (Cat. 631275). Briefly, 7 μg of the pLenti_Tet-coBxb1-2A-BFP_IRES-iCasp9-2A-Blast_rtTA3 vector was diluted in 600 μl of water, combined with the Lenti-X Packaging plasmid mix, and incubated for 10 minutes at room temperature. The resulting mixture was added dropwise to Lenti-X 293T cells cultured on a 10 cm dish and incubated at 37 °C with 5% CO₂. The culture medium was replaced the following day, and viral supernatants were harvested at 48 and 72 hours post-transfection and pooled. The collected supernatant was clarified by centrifugation at 300 × g for 5 minutes and passed through a 0.45 μm filter to remove cellular debris. The integrated landing pad construct encoded a doxycycline-inducible blue fluorescent protein (BFP) and a blasticidin resistance cassette, which were used to verify successful landing pad integration. The Hela cells plated on 6-well plate and each well contained a total volume of 2 mL, with virus-to-media ratios of 1.5 mL virus and 0.5 mL media, 1 mL virus and 1 mL media, 0.5 mL virus and 1.5 mL media, 250 µL virus and 1.85 mL media, or 100 µL virus and 1.9 mL media. After 24 hours, viral supernatant was removed and replaced with fresh culture medium containing doxycycline. BFP expression was evaluated 48 hours post-infection to identify cell populations infected at a multiplicity of infection (MOI) below 1. Following selection with 6 μg/ml blasticidin for one week, cells exhibiting the highest levels of BFP fluorescence were single-cell sorted into 96-well plates using a BD FACS Aria II (405 nm laser and a 450/50 nm emission filter). Following sorting, cells were cultured for 48 hours in doxycycline-containing medium only, after which blasticidin was reintroduced. Wells were monitored for two weeks and single-cell–derived colonies were identified and expanded in 6 well plates. Expanded clonal cell lines were subjected to dual-colour transfection using attB-eGFP and attB-mCherry constructs. A clone containing a single landing pad was identified by flow cytometry as cells expressing either eGFP or mCherry alone, with no dual-colour–positive population. The selected clone was expanded for downstream use.

### Library transfection

HeLa landing-pad cells were expanded in six T175 flasks and harvested 115 × 10⁶ cells for transfection. Cells were pelleted, washed once with PBS, and resuspended at 90 million cells in 740 µL Opti-MEM. 400 µg of library plasmid DNA was diluted in 160 µL water, and added to the cell solution to generate a final 900 µL transfection mixture. We used six R50*3 chambers and loaded each chamber with 50ul of the transfection mix and electroporated on MaxCyte GLA using default HeLa optimised program on the machine. Cells were allowed to recover for 20–30 min at 37 °C before transferring proportionally into 18 pre-warmed T75 flasks containing doxycycline. Cell viability was monitored following transfection. At 48 h post-transfection, an estimated 30–40% appeared dead and floating. Puromycin selection was initiated 72 h post-transfection using 3 µg/mL doxycycline and 0.4 µg/mL puromycin. After 48 h of selection, minimal cell death was observed, prompting concerns regarding puromycin efficacy. Cells were therefore detached, redistributed into nine T225 flasks, and puromycin concentration was increased to 0.8 µg/mL. After an additional 48 h, extensive cell death (∼30–40%) was observed, with cultures at ∼50% confluency. Based on cumulative losses, approximately 15% of the originally transfected population remained, corresponding to ∼13.5 million cells, which was considered sufficient library representation. Medium changed every two to three days. Once the flasks were confluent, cells were harvested yielding 252 million cells, resuspended in 30 mL freezing medium and stored as 30 cryovials in liquid nitrogen.

### Genomic DNA extraction, barcode amplification and sequencing

Cells were gated for singlets and sorted on a BD FACS Aria II into four populations according to fluorescence intensity in the 640–670/14 A channel. At least 10 million cells were collected per bin. Following sorting, cells from each bin were aliquoted into four 1.5 mL tubes, pelleted by centrifugation, and stored at −20 °C prior to genomic DNA extraction and library preparation. Genomic DNA from cells in each sorted bin was isolated using phenol–chloroform extraction. Cell pellets were resuspended in 500 μL of lysis buffer containing 100 mM Tris (pH 8.5), 0.5 mM EDTA, 0.2% SDS, 200 mM NaCl, and 25 μg RNase A, and incubated at 37°C for 1 hour. Proteinase K (700 μg; New England Biolabs, P8107S) was then added, followed by overnight incubation at 56°C. An equal volume of phenol:chloroform:isoamyl alcohol (25:24:1) was added, and samples were vortexed vigorously and centrifuged at high speed for 5 minutes. The aqueous phase was transferred to a fresh 1.5 mL tube and subjected to a second extraction with an equal volume of phenol:chloroform:isoamyl alcohol, followed by vigorous vortexing and centrifugation for 5 minutes. The aqueous layer was transferred again to a new tube, and DNA was precipitated with 1mL ethanol and resuspended in 100 μL of nuclease-free water. The four tubes corresponding to each bin were subsequently pooled into a single tube containing a total of 400 μL of extracted genomic DNA. Genomic DNA barcodes were amplified by PCR. To prevent amplification of barcodes derived from any residual extragenomic barcoded pMV2-Parkin plasmid, the reverse primer was designed to anneal to a genomic DNA region located downstream of the plasmid integration site. For each sorted bin, purified genomic DNA was divided among multiple 25 μL PCR reactions. Each reaction contained 12.5 μL of Phusion High-Fidelity PCR Master Mix with HF buffer (New England Biolabs, M0531S), 500 ng of genomic DNA, 10 μM of a reverse primer carrying the Illumina P5 sequence (VA04), and 10 μM of an indexed forward primer incorporating the Illumina P7 sequence (VA05) that anneals upstream of the barcode. PCR cycling conditions consisted of an initial denaturation at 96°C for 2 minutes, followed by 24 cycles of denaturation at 96°C for 30 seconds, annealing at 65°C for 30 seconds, and extension at 72°C for 40 seconds, with a final extension at 72 °C for 1 minute. PCR products from each bin were pooled, and 300 μL of the combined material was purified using the MinElute PCR Purification Kit and eluted in 50 μL of water. The purified samples were then sent to WTCRF for size selection of the 188 bp PCR product with equimolar mixing and sequencing on an Illumina NextSeq platform using a custom sequencing primer (VA06). Demultiplexed sequencing reads were processed with Trimmomatic to retain only the 30-nucleotide barcode region. These trimmed sequences were subsequently aligned against the barcode library defined by PacBio sequencing using Enrich2 software, generating counts for each library barcode in every FACS sorting bin. Raw Illumina sequencing data and processed files have been deposited in the Gene Expression Omnibus (GEO) under accession XXXX. Analysis scripts are publicly available at https://zenodo.org/records/XXXX.

### Calculation of variant function scores and false discovery rates

After counting the barcodes in each bin with Enrich2 (Rubin et al., 2017) we then used a previously published method for calculation of variant function scores and false discovery rates [57]. For FDR estimation based on stop-codon barcodes, we used the same framework but bootstrapped stop-codon barcodes instead of wild-type barcodes. Parkin variant effect predictor scores were from [69].

### Computation of biallelic function scores

The following filtering steps were applied to MDSGene data in order to calculate biallelic functional scores and correlate with the age at disease onset:

1. Read 2934 patient info from MDSGene data;
2. Keep only patients with at least one missense mutations in one allele (either in mut1 column (allele 1) or mut2 column (allele 2)): left 867
3. Remove digenic genotypes: left 856
4. Remove patients with unknown genotype info, represented with -99: left 853
5. Remove patients with hom genotype but mut1 is structural variation and mut2 is missense (ambiguous cases): left 851
6. Remove missense mutations not present in our data, wrong annotations (I298S, M1T, R247G, M1L, A469T, E309*, A339A, M318L, A17D, G234W): left 823.
7. Remove M1L and M1T as we didn’t mutate them, also removed others like A17D, the position 17 is not A, this is the case for the others as pie_with_external_labels
8. Remove patients with unknown aao info represented with -99 or NaN: left 634
9. Remove ambiguous patients_genotype, just one case represented with "compound het (hom)": left 633

### Generating single substitution-stable cell lines

Primers (**Supplementary Table 3**) were designed to introduce R6F, G12E, R42E, V70T, V186H, E322H, Q326E, and W403A substitutions into the attB_PRKN_IRES_PuroR (pMV2) plasmid using the Q5 Site-Directed Mutagenesis Kit (New England Biolabs, E0554S) following manufacturer’s instructions. For the R42P, S65A, R275W, and C431F, variants were subcloned from pcDNA5D-FRT/TO/PRKN constructs obtained from the MRC PPU Reagents and Services, University of Dundee (R42P, DU43232; S65A, DU23315; R275W, DU23337; C431F, DU23340). Inserts were amplified by PCR using VA01/VA02 primers, followed by digestion with KasI and EcoRI. The resulting fragments were purified, cloned into the Gateway entry vector pAINt, and subsequently transferred into attB_PRKN_IRES_PuroR (pMV2) by Gateway LR reaction. All constructs were transformed into NEB^Ⓡ^ 5-alpha *E. coli* and sequence-verified by whole-plasmid sequencing. For making stable cell lines expressing the single variants, 500 ng of each plasmid was transfected into 3×10⁵ Bxb1 landing-pad HeLa cells using FuGENE^Ⓡ^ 6 Transfection Reagent (Promega, E2691) on a 12-well plate followed by doxycycline (3 µg/mL) induction and puromycin selection (1 µg/mL) 14 days prior to flow cytometry.

## Acknowledgements

This work was funded by the Michael J. Fox Foundation for Parkinson’s Research (MJFF) through grant MJFF-024896 to G.K., M.M.K.M., E.M.S. and D.R.A. M.M.K.M. is supported by Parkinson’s UK and the UK Dementia Research Institute through UK DRI Ltd, principally funded by the Medical Research Council. G.K and M.C. are supported by the Medical Research Council (MRC) Human Genetics Unit core grant MC_UU_00035/8. L.R. is supported by a Medical Research Council PhD studentship.

## Figure legends

**Supplementary Figure 1.**
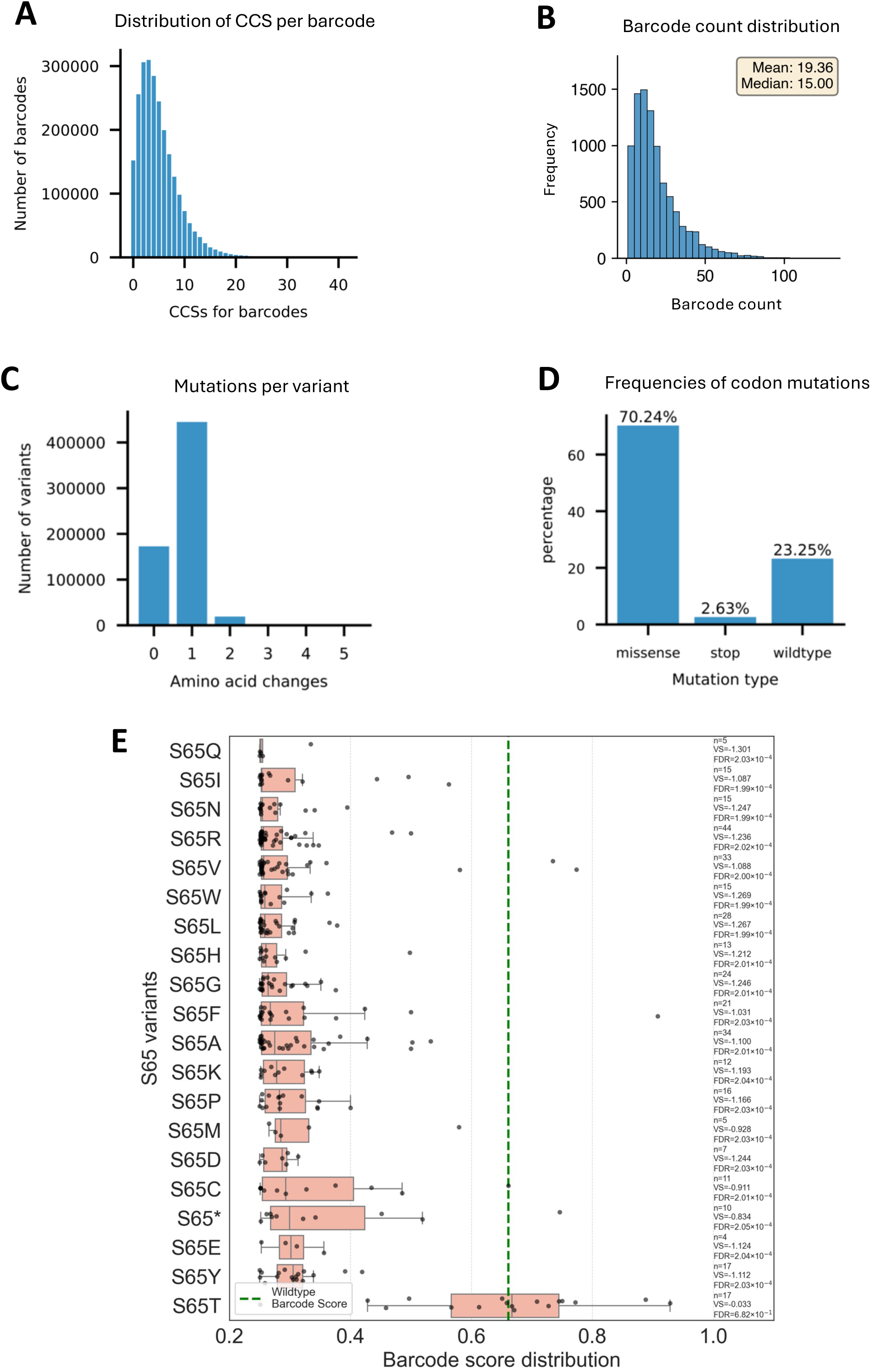
Composition of Parkin library and barcode scores of selected variants. **A)** Number of circular consensus sequences (CCS) in the PacBio analysis used for barcode-variant phasing. **B)** Distribution of numbers of barcodes per variant. **C)** Numbers of amino acid changes per variant. **D)** Frequencies of missense, synonymous and nonsense variants. **E)** Raw barcode scores of all S65 variants in the pS65-Ub assay.

**Supplementary Figure 2.**
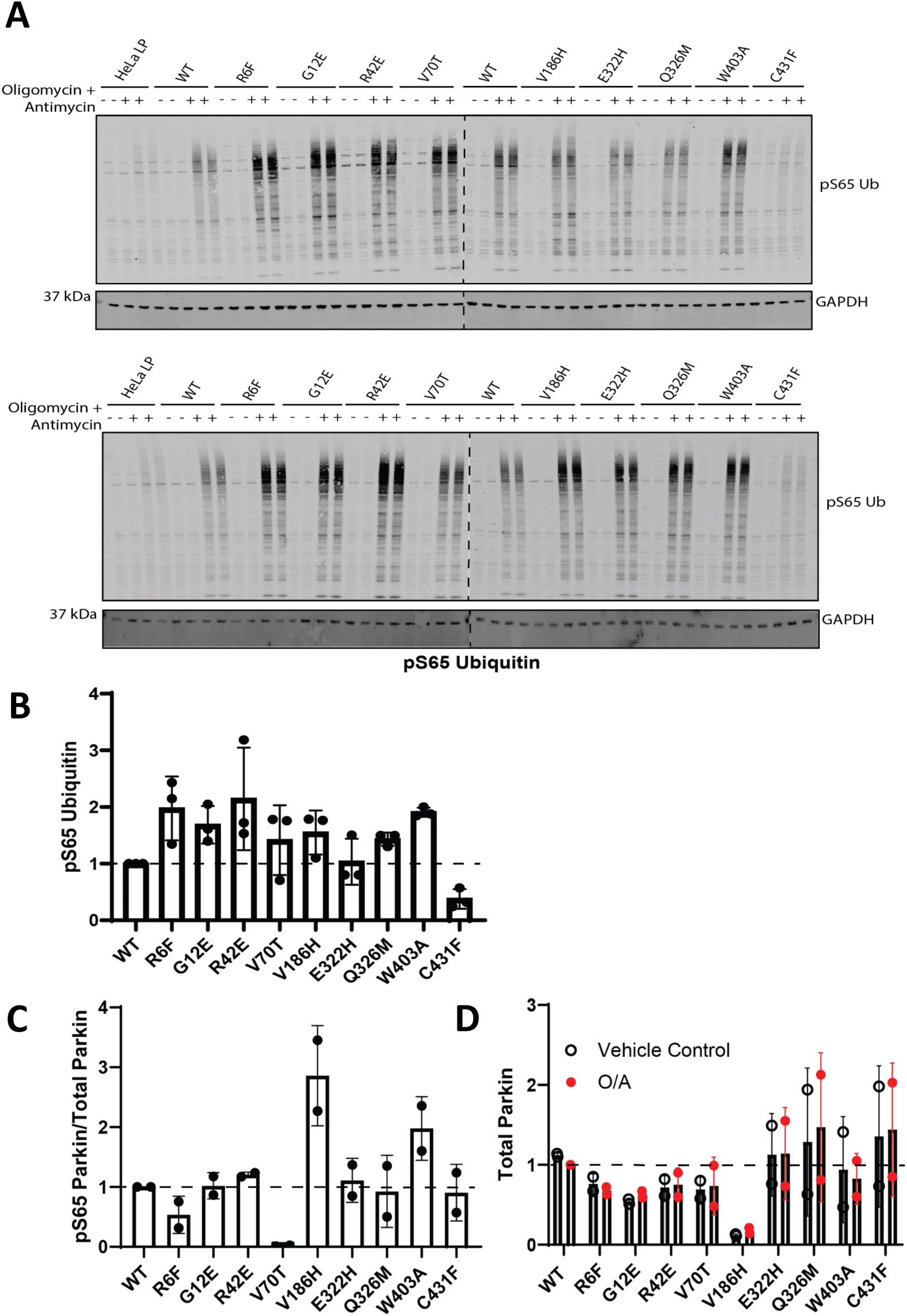
Western blot analysis of cells expressing selected Parkin variants. **A)** pS65-Ub analysis performed in two biological replicates in addition to that shown in Fig. 2. Each biological replicate was performed on a different day and comprises a pair of technical replicates for each condition. (-) and (+) represent untreated cells and cells treated with 1 μM Oligomycin and 10 μM Antimycin for 45 minutes, respectively. **B)** Quantification of data in panel A. **C and D)** Quantification of Parkin and pS65 Parkin immunoblots from Figure 2.

**Supplementary Figure 3.**
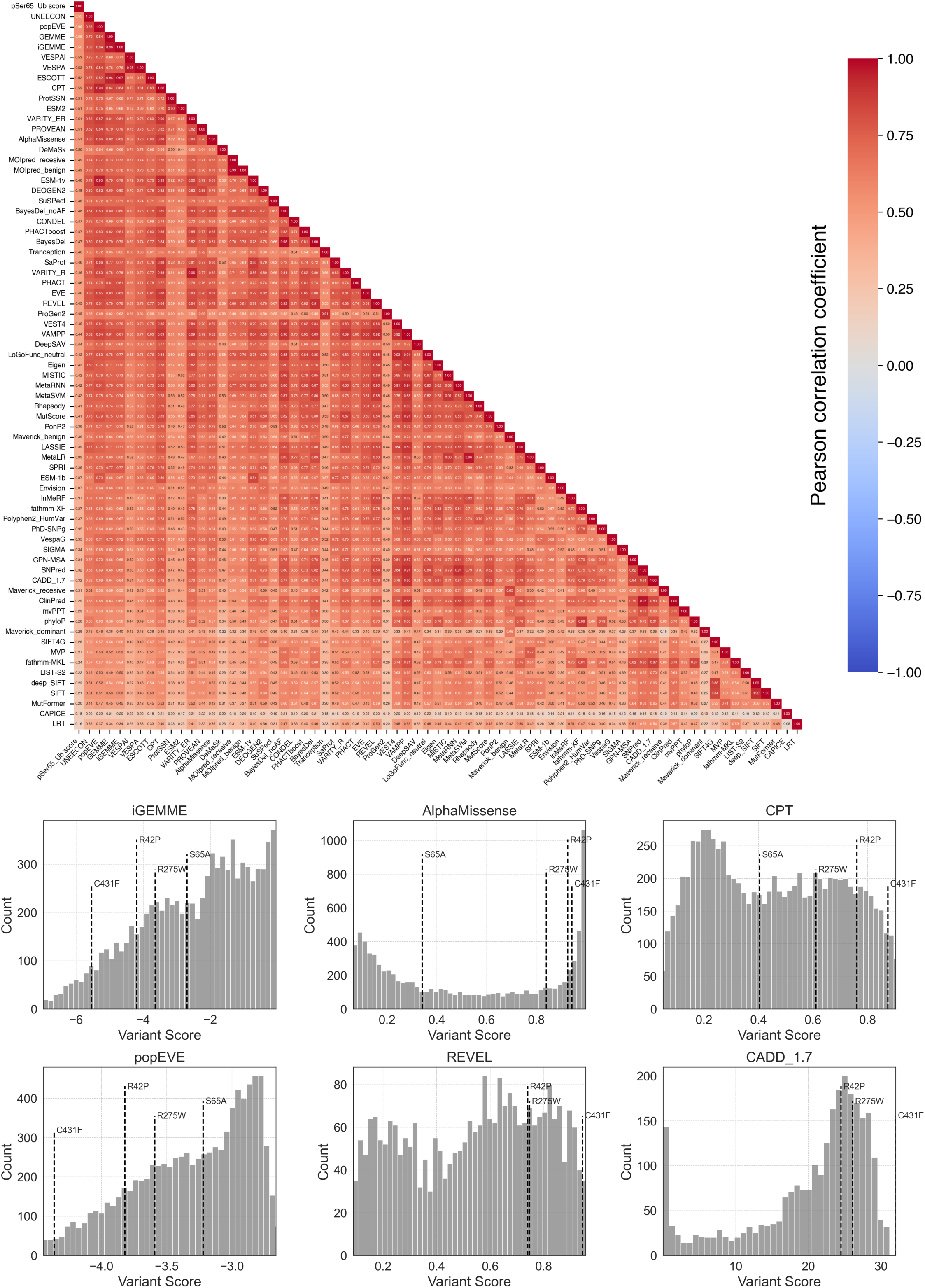
Comparison of pS65-Ub functional scores with scores from variant effect predictors (VEPs). **A)** Pearson correlation matrix between pS65-Ub functional scores and scores from 70 VEPs. VEP scores were inverted where necessary to match the orientation of variant scores in the data (lower scores indicating lower function and higher pathogenicity). **B)** Distribution of missense variant scores for selected VEPs, with selected variants indicated by dashed lines.

**Supplementary Figure 4.**
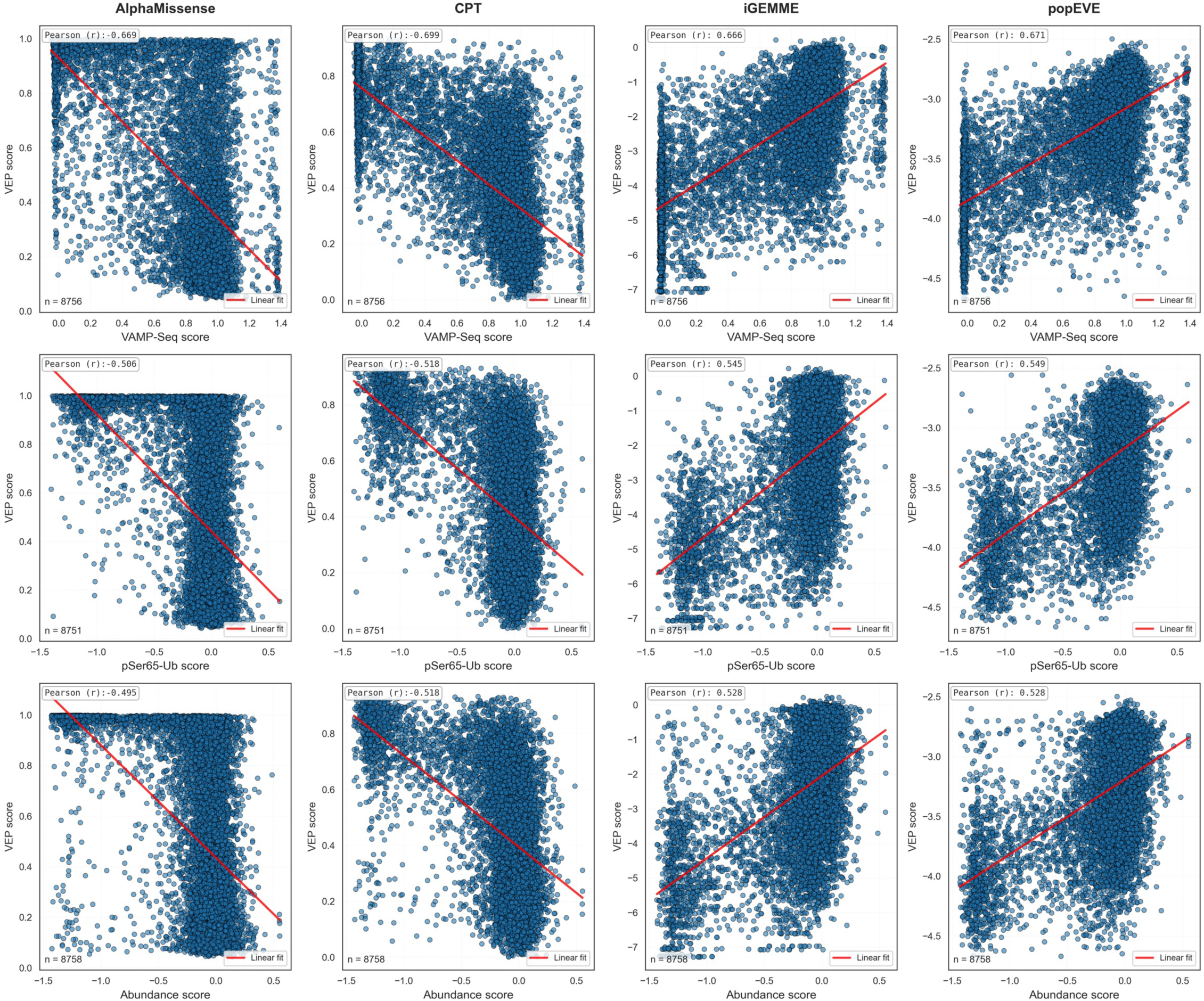
Comparison of experimental and computational Parkin variant effect scores. VAMP-Seq scores from Clausen et al. (2024, top), pS65-Ub functional scores (middle), and abundance scores from the present study (bottom), are correlated against AlphaMissense, CPT, iGEMME, and popEVE pathogenicity scores.

**Supplementary Figure 5.**
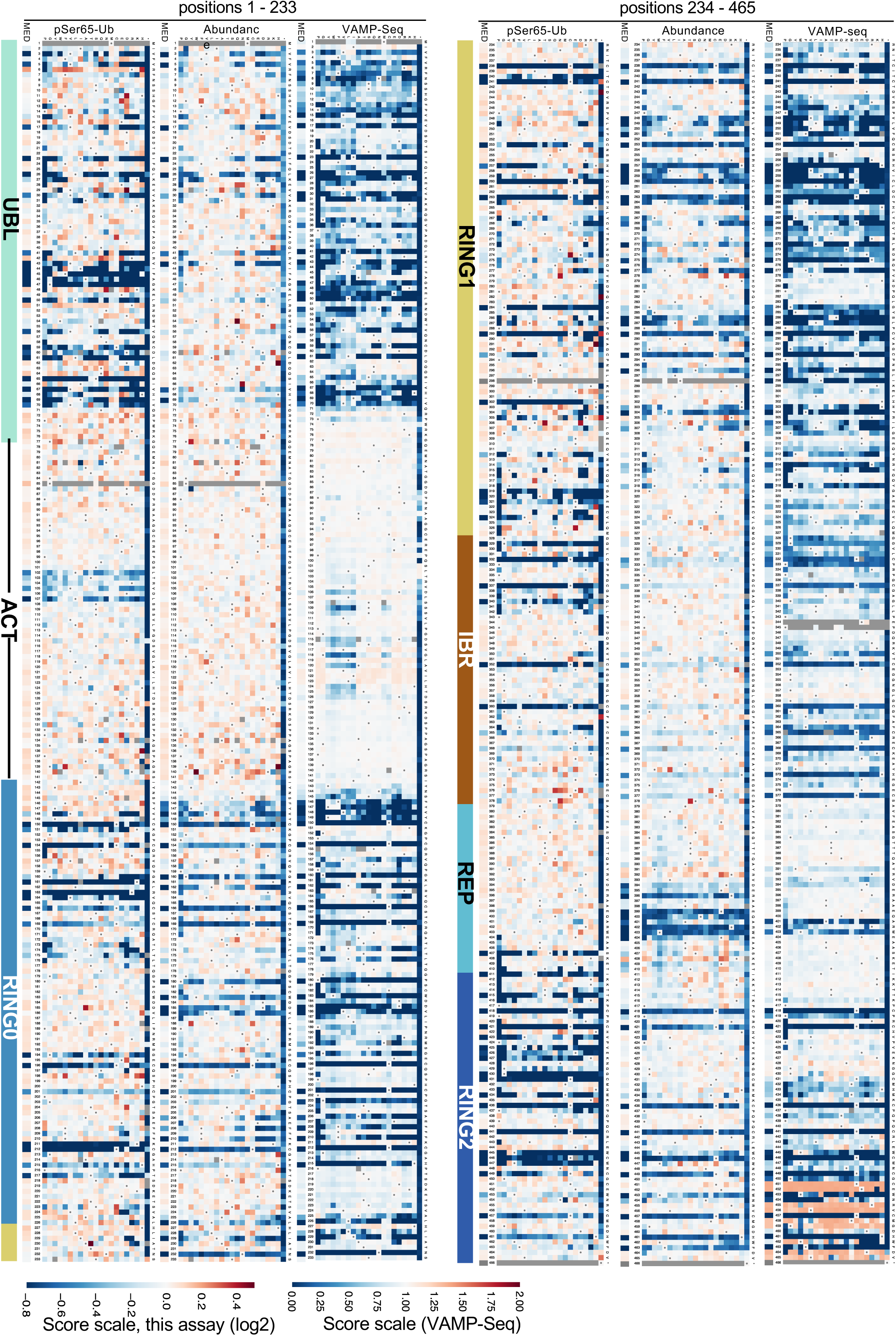
Comparison of mutational effects on Parkin abundance and function. pS65-Ub functional scores (left), abundance scores from the present study (middle), and VAMP-Seq scores from Clausen et al. (2024, right) are shown alongside each other as heatmaps, with colours as in Figure 2.

**Supplementary Figure 6.**
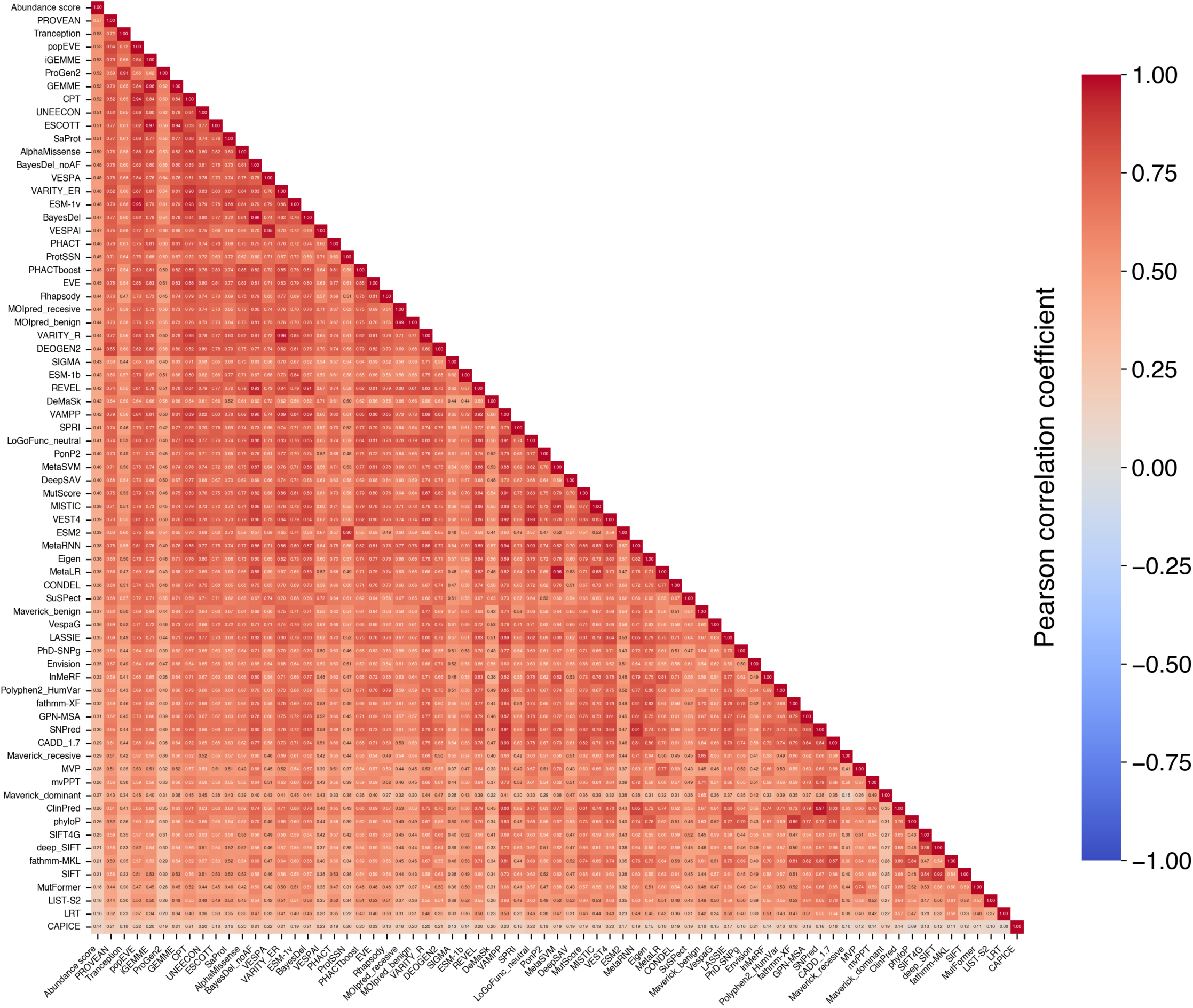
Pearson correlation matrix between Parkin abundance scores from this study and scores from 70 VEPs.

**Supplementary Figure 7.**
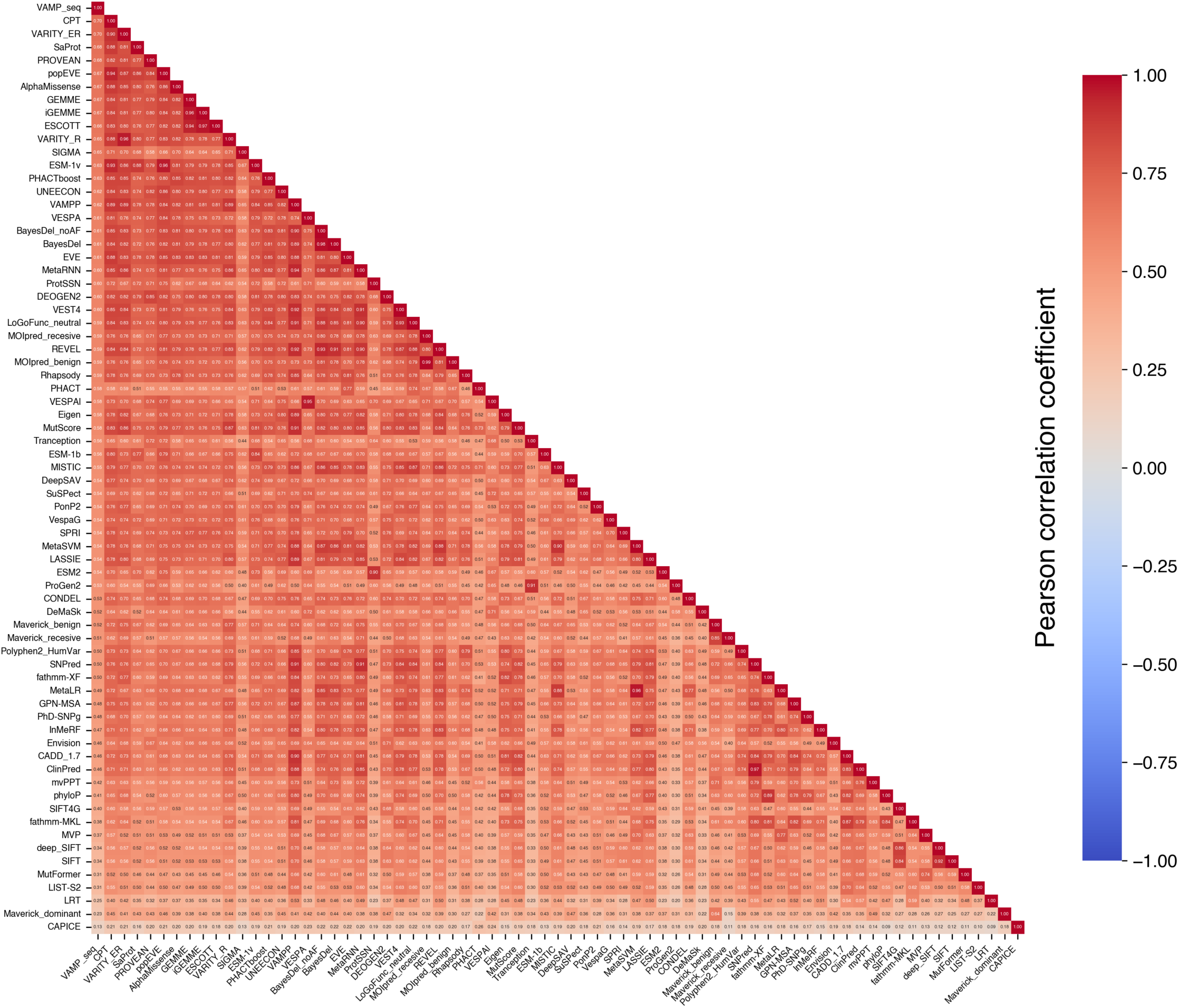
Pearson correlation matrix between Parkin VAMP-Seq scores from Clausen et al. (2024) and scores from 70 VEPs.

**Supplementary Figure 8.**
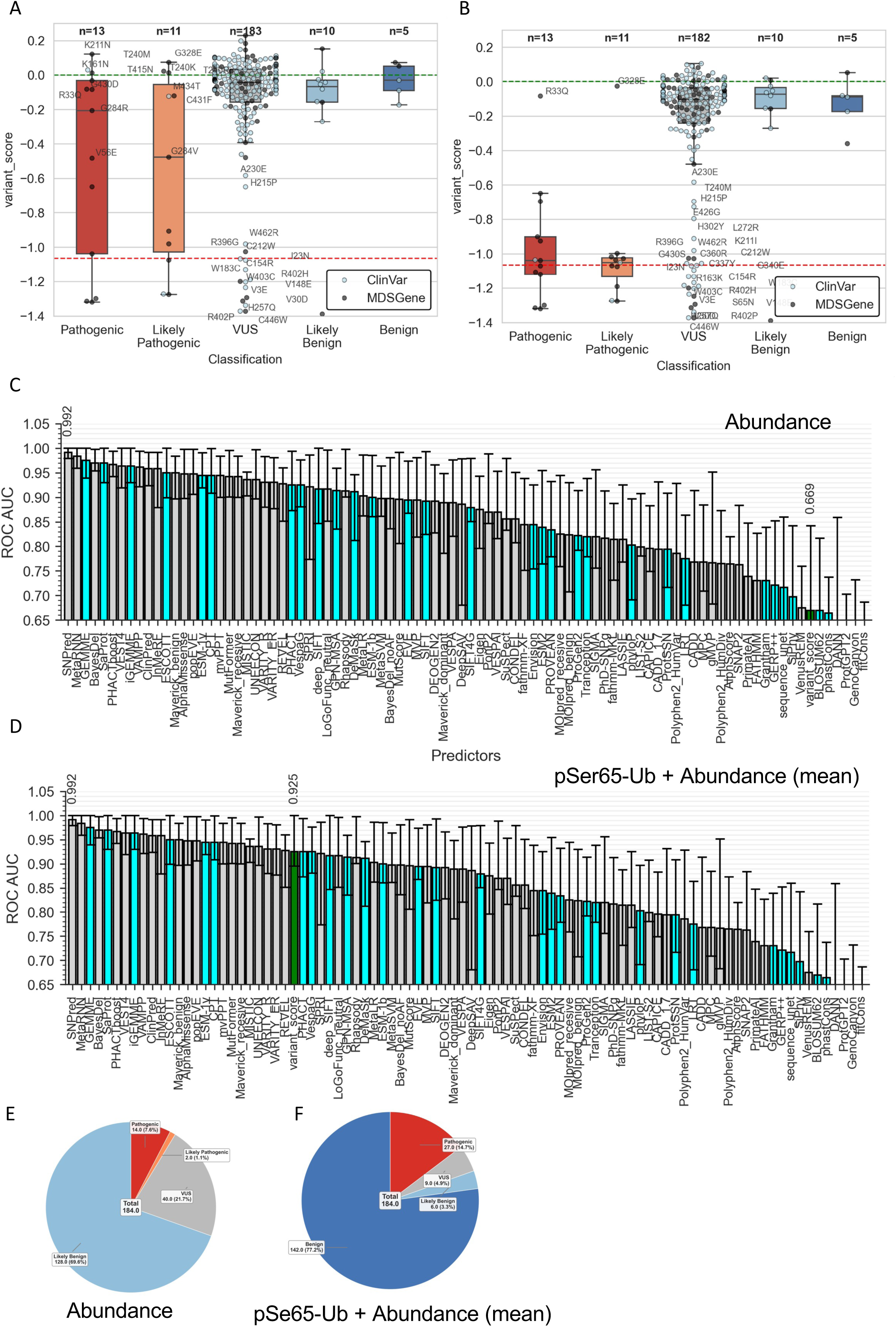
Comparison of Parkin variant abundance scores with clinical annotations. **A)** Boxplots showing abundance scores for variants stratified according to their clinical annotations as in Fig. 3. **B)** Boxplots showing the minimum of pS65-Ub and abundance scores for variants stratified as above. **C and D)** Distribution of ROC AUC scores in the classification of benign vs pathogenic variants for VEPs and MAVE abundance scores (C) or means of pS65-Ub and abundance scores (D). **E) and F)** Classification of VUS using their abundance scores (E) or means of pS65-Ub and abundance scores (F).

**Supplementary Figure 9.**
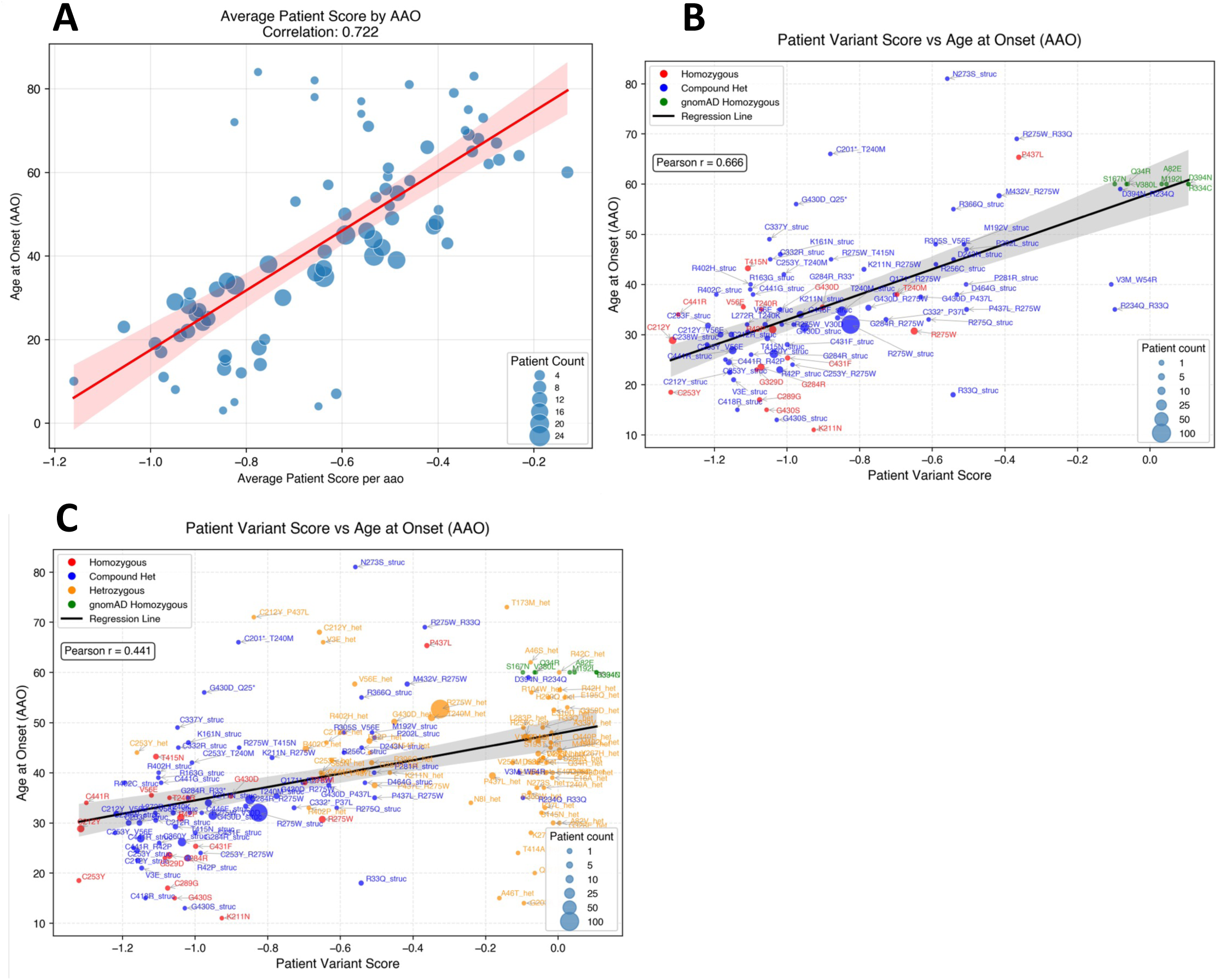
Prediction of PD age at onset (AAO) based on patient-level biallelic scores. **A-C)** As in Figure 4, but using biallelic scores derived from the minimum of pS65-Ub and abundance scores.

**Supplementary Figure 10.**
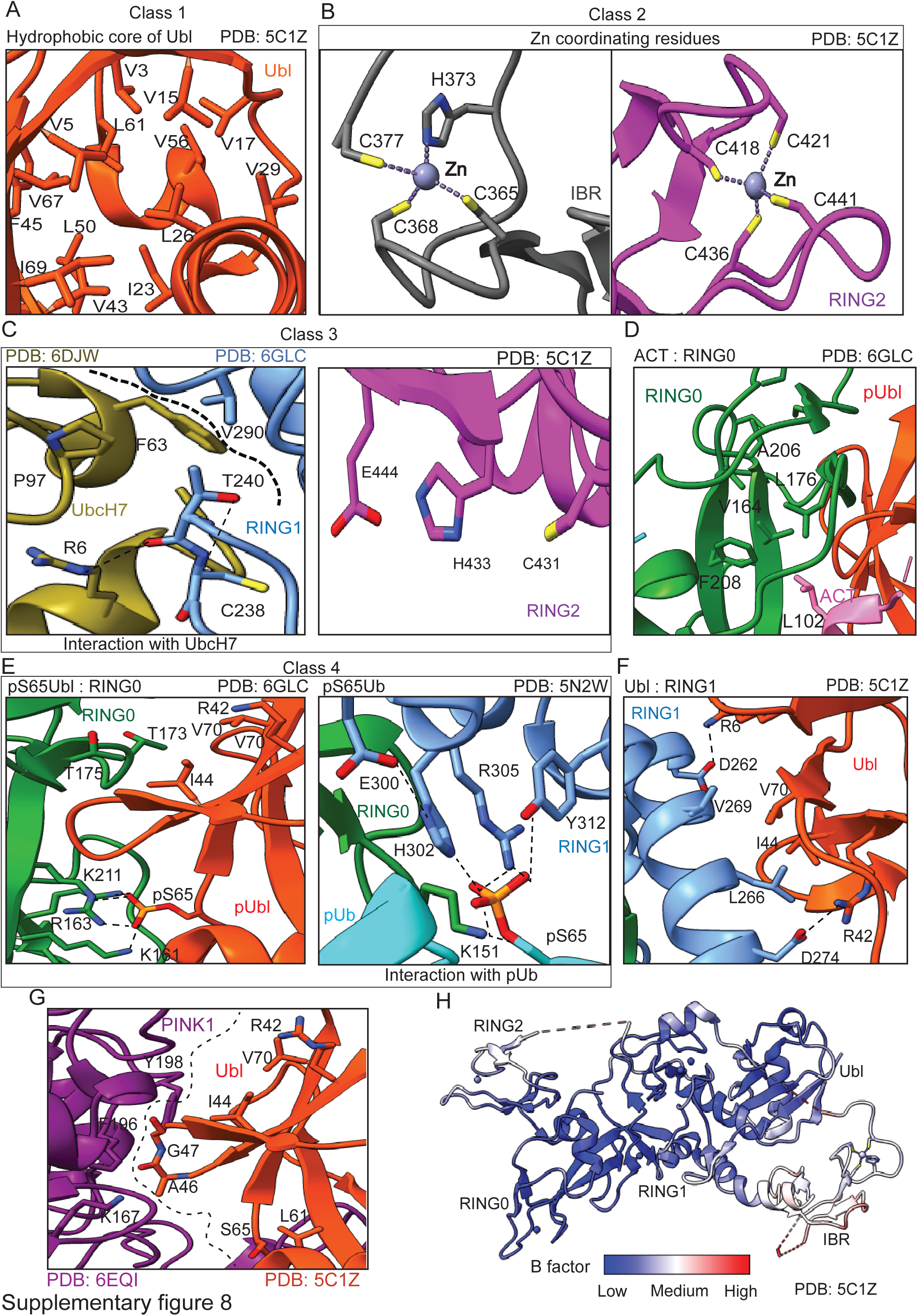
Interaction interfaces of Parkin variants of interest (VOI) mapped onto the Parkin structure A-E. **(A–E)** Detailed interaction interfaces of selected inactivating Parkin variants from different mechanistic classes, as shown in Fig. 6A. **(F)** Interaction interface between the UBL and RING1 domains, highlighting activating mutations located in the UBL domain, as shown in Fig. 6A. **(G)** Interaction interface between the Parkin UBL domain and PINK1, highlighting key Parkin residues that may influence Parkin phosphorylation. The Parkin UBL domain was modelled onto PINK1 by superimposition onto ubiquitin in the PINK1–Ub crystal structure (PDB ID: 6EQI). **(H)** B-factor analysis of the Parkin crystal structure (PDB ID: 5C1Z).

## Notes

https://parkin-app-umfr.onrender.com/

### Competing Interest Statement

The authors have declared no competing interest.

## References

1. Goedert M, Spillantini MG, Del Tredici K, Braak H: 100 years of Lewy pathology. Nat Rev Neurol 2013, 9:13–24.

2. Bloem BR, Okun MS, Klein C: Parkinson’s disease. Lancet 2021, 397:2284–2303.

3. Lim SY, Klein C: Parkinson’s Disease is Predominantly a Genetic Disease. J Parkinsons Dis 2024, 14:467–482.

4. Dorsey ER, Bloem BR: Parkinson’s Disease Is Predominantly an Environmental Disease. J Parkinsons Dis 2024, 14:451–465.

5. Skrahina V, Gaber H, Vollstedt EJ, Forster TM, Usnich T, Curado F, Bruggemann N, Paul J, Bogdanovic X, Zulbahar S, et al: The Rostock International Parkinson’s Disease (ROPAD) Study: Protocol and Initial Findings. Mov Disord 2021, 36:1005–1010.

6. Cook L, Verbrugge J, Schwantes-An TH, Schulze J, Foroud T, Hall A, Marder KS, Mata IF, Mencacci NE, Nance MA, et al: Parkinson’s disease variant detection and disclosure: PD GENEration, a North American study. Brain 2024, 147:2668–2679.

7. Lucking CB, Durr A, Bonifati V, Vaughan J, De Michele G, Gasser T, Harhangi BS, Meco G, Denefle P, Wood NW, et al: Association between early-onset Parkinson’s disease and mutations in the parkin gene. N Engl J Med 2000, 342:1560–1567.

8. Harper JW, Ordureau A, Heo JM: Building and decoding ubiquitin chains for mitophagy. Nat Rev Mol Cell Biol 2018, 19:93–108.

9. Antico O, Thompson PW, Hertz NT, Muqit MMK, Parton LE: Targeting mitophagy in neurodegenerative diseases. Nat Rev Drug Discov 2025, 24:276–299.

10. Trempe JF, Gehring K: Structural Mechanisms of Mitochondrial Quality Control Mediated by PINK1 and Parkin. J Mol Biol 2023, 435:168090.

11. Gundogdu M, Tadayon R, Salzano G, Shaw GS, Walden H: A mechanistic review of Parkin activation. Biochim Biophys Acta Gen Subj 2021, 1865:129894.

12. Sherawat M, Kumar A, Lenka DR, Kumar A: Cis or trans: a puzzle of Parkin activation mechanism. Essays Biochem 2026, 69.

13. Kazlauskaite A, Kondapalli C, Gourlay R, Campbell DG, Ritorto MS, Hofmann K, Alessi DR, Knebel A, Trost M, Muqit MM: Parkin is activated by PINK1-dependent phosphorylation of ubiquitin at Ser65. Biochem J 2014, 460:127–139.

14. Kane LA, Lazarou M, Fogel AI, Li Y, Yamano K, Sarraf SA, Banerjee S, Youle RJ: PINK1 phosphorylates ubiquitin to activate Parkin E3 ubiquitin ligase activity. J Cell Biol 2014, 205:143–153.

15. Koyano F, Okatsu K, Kosako H, Tamura Y, Go E, Kimura M, Kimura Y, Tsuchiya H, Yoshihara H, Hirokawa T, et al: Ubiquitin is phosphorylated by PINK1 to activate parkin. Nature 2014, 510:162–166.

16. Kondapalli C, Kazlauskaite A, Zhang N, Woodroof HI, Campbell DG, Gourlay R, Burchell L, Walden H, Macartney TJ, Deak M, et al: PINK1 is activated by mitochondrial membrane potential depolarization and stimulates Parkin E3 ligase activity by phosphorylating Serine 65. Open Biol 2012, 2:120080.

17. Ordureau A, Sarraf SA, Duda DM, Heo JM, Jedrychowski MP, Sviderskiy VO, Olszewski JL, Koerber JT, Xie T, Beausoleil SA, et al: Quantitative proteomics reveal a feedforward mechanism for mitochondrial PARKIN translocation and ubiquitin chain synthesis. Mol Cell 2014, 56:360–375.

18. Ordureau A, Heo JM, Duda DM, Paulo JA, Olszewski JL, Yanishevski D, Rinehart J, Schulman BA, Harper JW: Defining roles of PARKIN and ubiquitin phosphorylation by PINK1 in mitochondrial quality control using a ubiquitin replacement strategy. Proc Natl Acad Sci U S A 2015, 112:6637–6642.

19. Oliveira SA, Scott WK, Martin ER, Nance MA, Watts RL, Hubble JP, Koller WC, Pahwa R, Stern MB, Hiner BC, et al: Parkin mutations and susceptibility alleles in late-onset Parkinson’s disease. Ann Neurol 2003, 53:624–629.

20. Klein C, Lohmann-Hedrich K, Rogaeva E, Schlossmacher MG, Lang AE: Deciphering the role of heterozygous mutations in genes associated with parkinsonism. Lancet Neurol 2007, 6:652–662.

21. Hedrich K, Eskelson C, Wilmot B, Marder K, Harris J, Garrels J, Meija-Santana H, Vieregge P, Jacobs H, Bressman SB, et al: Distribution, type, and origin of Parkin mutations: review and case studies. Mov Disord 2004, 19:1146–1157.

22. Kasten M, Hartmann C, Hampf J, Schaake S, Westenberger A, Vollstedt EJ, Balck A, Domingo A, Vulinovic F, Dulovic M, et al: Genotype-Phenotype Relations for the Parkinson’s Disease Genes Parkin, PINK1, DJ1: MDSGene Systematic Review. Mov Disord 2018, 33:730–741.

23. Chaugule VK, Burchell L, Barber KR, Sidhu A, Leslie SJ, Shaw GS, Walden H: Autoregulation of Parkin activity through its ubiquitin-like domain. EMBO J 2011, 30:2853–2867.

24. Trempe JF, Sauve V, Grenier K, Seirafi M, Tang MY, Menade M, Al-Abdul-Wahid S, Krett J, Wong K, Kozlov G, et al: Structure of parkin reveals mechanisms for ubiquitin ligase activation. Science 2013, 340:1451–1455.

25. Yi W, MacDougall EJ, Tang MY, Krahn AI, Gan-Or Z, Trempe JF, Fon EA: The landscape of Parkin variants reveals pathogenic mechanisms and therapeutic targets in Parkinson’s disease. Hum Mol Genet 2019, 28:2811–2825.

26. Stevens MU, Croteau N, Eldeeb MA, Antico O, Zeng ZW, Toth R, Durcan TM, Springer W, Fon EA, Muqit MM, Trempe JF: Structure-based design and characterization of Parkin-activating mutations. Life Sci Alliance 2023, 6.

27. Traynor R, Moran J, Stevens M, Antico O, Knebel A, Behrouz B, Merchant K, Hastie CJ, Davies P, Muqit MMK, De Cesare V: Design and high-throughput implementation of MALDI-TOF/MS-based assays for Parkin E3 ligase activity. Cell Rep Methods 2024, 4:100712.

28. Clausen L, Voutsinos V, Cagiada M, Johansson KE, Gronbaek-Thygesen M, Nariya S, Powell RL, Have MKN, Oestergaard VH, Stein A, et al: A mutational atlas for Parkin proteostasis. Nat Commun 2024, 15:1541.

29. Lange LM, Fang ZH, Makarious MB, Kuznetsov N, Brolin KA, Ballard S, Bardien S, Doquenia ML, Heutink P, Houlden H, et al: The Global Landscape of Genetic Variation in Parkinson’s disease: Multi-Ancestry Insights into Established Disease Genes and their Translational Relevance. medRxiv 2025.

30. Çubuk H, Plech, M., Aslanzadeh, V., Zikanova, M., Skopova, V., Kmoch, S., Shen, Y., Marsh, J.A., Kudla, G.: Mechanistic Modelling of Recessive Disease through Allelic Integration of Variant Effects. bioRxiv 2025.

31. Wrenbeck EE, Klesmith JR, Stapleton JA, Adeniran A, Tyo KE, Whitehead TA: Plasmid-based one-pot saturation mutagenesis. Nat Methods 2016, 13:928–930.

32. Hung CM, Lombardo PS, Malik N, Brun SN, Hellberg K, Van Nostrand JL, Garcia D, Baumgart J, Diffenderfer K, Asara JM, Shaw RJ: AMPK/ULK1-mediated phosphorylation of Parkin ACT domain mediates an early step in mitophagy. Sci Adv 2021, 7.

33. Cookson MR, Lockhart PJ, McLendon C, O’Farrell C, Schlossmacher M, Farrer MJ: RING finger 1 mutations in Parkin produce altered localization of the protein. Hum Mol Genet 2003, 12:2957–2965.

34. Fiesel FC, Caulfield TR, Moussaud-Lamodiere EL, Ogaki K, Dourado DF, Flores SC, Ross OA, Springer W: Structural and Functional Impact of Parkinson Disease-Associated Mutations in the E3 Ubiquitin Ligase Parkin. Hum Mutat 2015, 36:774–786.

35. Gladkova C, Maslen SL, Skehel JM, Komander D: Mechanism of parkin activation by PINK1. Nature 2018, 559:410–414.

36. Fakih R, Sauve V, Gehring K: Structure of the second phosphoubiquitin-binding site in parkin. J Biol Chem 2022, 298:102114.

37. Lenka DR, Dahe SV, Antico O, Sahoo P, Prescott AR, Muqit MMK, Kumar A: Additional feedforward mechanism of Parkin activation via binding of phospho-UBL and RING0 in trans. Elife 2024, 13.

38. Huq TS, Luo J, Fakih R, Sauve V, Gehring K: Naturally occurring hyperactive variants of human parkin. Commun Biol 2024, 7:961.

39. Beasley SA, Hristova VA, Shaw GS: Structure of the Parkin in-between-ring domain provides insights for E3-ligase dysfunction in autosomal recessive Parkinson’s disease. Proc Natl Acad Sci U S A 2007, 104:3095–3100.

40. Kazlauskaite A, Kelly V, Johnson C, Baillie C, Hastie CJ, Peggie M, Macartney T, Woodroof HI, Alessi DR, Pedrioli PG, Muqit MM: Phosphorylation of Parkin at Serine65 is essential for activation: elaboration of a Miro1 substrate-based assay of Parkin E3 ligase activity. Open Biol 2014, 4:130213.

41. Seirafi M, Kozlov G, Gehring K: Parkin structure and function. FEBS J 2015, 282:2076–2088.

42. McWilliams TG, Barini E, Pohjolan-Pirhonen R, Brooks SP, Singh F, Burel S, Balk K, Kumar A, Montava-Garriga L, Prescott AR, et al: Phosphorylation of Parkin at serine 65 is essential for its activation in vivo. Open Biol 2018, 8.

43. Badonyi M, Marsh JA: acmgscaler: an R package and Colab for standardized gene-level variant effect score calibration within the ACMG/AMP framework. Bioinformatics 2025, 41.

44. Livesey BJ, Marsh JA: Updated benchmarking of variant effect predictors using deep mutational scanning. Mol Syst Biol 2023, 19:e11474.

45. Wauer T, Simicek M, Schubert A, Komander D: Mechanism of phospho-ubiquitin-induced PARKIN activation. Nature 2015, 524:370–374.

46. Kumar A, Aguirre JD, Condos TE, Martinez-Torres RJ, Chaugule VK, Toth R, Sundaramoorthy R, Mercier P, Knebel A, Spratt DE, et al: Disruption of the autoinhibited state primes the E3 ligase parkin for activation and catalysis. EMBO J 2015, 34:2506–2521.

47. Schubert AF, Gladkova C, Pardon E, Wagstaff JL, Freund SMV, Steyaert J, Maslen SL, Komander D: Structure of PINK1 in complex with its substrate ubiquitin. Nature 2017, 552:51–56.

48. Clark IE, Dodson MW, Jiang C, Cao JH, Huh JR, Seol JH, Yoo SJ, Hay BA, Guo M: Drosophila pink1 is required for mitochondrial function and interacts genetically with parkin. Nature 2006, 441:1162–1166.

49. Palacino JJ, Sagi D, Goldberg MS, Krauss S, Motz C, Wacker M, Klose J, Shen J: Mitochondrial dysfunction and oxidative damage in parkin-deficient mice. J Biol Chem 2004, 279:18614–18622.

50. Narendra D, Tanaka A, Suen DF, Youle RJ: Parkin is recruited selectively to impaired mitochondria and promotes their autophagy. J Cell Biol 2008, 183:795–803.

51. Narendra DP, Jin SM, Tanaka A, Suen DF, Gautier CA, Shen J, Cookson MR, Youle RJ: PINK1 is selectively stabilized on impaired mitochondria to activate Parkin. PLoS Biol 2010, 8:e1000298.

52. Ordureau A, Paulo JA, Zhang J, An H, Swatek KN, Cannon JR, Wan Q, Komander D, Harper JW: Global Landscape and Dynamics of Parkin and USP30-Dependent Ubiquitylomes in iNeurons during Mitophagic Signaling. Mol Cell 2020, 77:1124–1142 e1110.

53. Terreni L, Calabrese E, Calella AM, Forloni G, Mariani C: New mutation (R42P) of the parkin gene in the ubiquitinlike domain associated with parkinsonism. Neurology 2001, 56:463–466.

54. Safadi SS, Shaw GS: A disease state mutation unfolds the parkin ubiquitin-like domain. Biochemistry 2007, 46:14162–14169.

55. Sigmarsdóttir ES, Vasileios, V., Johansson, K.E., Henrichs, I.K., Buhrmann, A.,, Lindorff-Larsen K, Rasmus Hartmann-Petersen, R.: Comprehensive Variant Effect Map of Parkin-Mediated Mitophagy in Parkinson’s Disease. BioRxiv 2026.

56. McDonnell AF, Plech M, Livesey BJ, Gerasimavicius L, Owen LJ, Hall HN, FitzPatrick DR, Marsh JA, Kudla G: Deep mutational scanning quantifies DNA binding and predicts clinical outcomes of PAX6 variants. BioRxiv 2023.

57. Aslanzadeh V, Brierley GV, Kumar R, Cubuk H, Vigouroux C, Matreyek KA, Kudla G, Semple RK: Deep mutational scanning of the human insulin receptor ectodomain to inform precision therapy for insulin resistance. Nat Commun 2025, 16:9143.

58. Gersing S, Cagiada M, Gebbia M, Gjesing AP, Cote AG, Seesankar G, Li R, Tabet D, Weile J, Stein A, et al: A comprehensive map of human glucokinase variant activity. Genome Biol 2023, 24:97.

59. Garcia EM, Lue NZ, Liang JK, Lieberman WK, Hwang DD, Woods JC, Liau BB: Base Editor Scanning Reveals Activating Mutations of DNMT3A. ACS Chem Biol 2023, 18:2030–2038.

60. Kumar A, Chaugule VK, Condos TEC, Barber KR, Johnson C, Toth R, Sundaramoorthy R, Knebel A, Shaw GS, Walden H: Parkin-phosphoubiquitin complex reveals cryptic ubiquitin-binding site required for RBR ligase activity. Nat Struct Mol Biol 2017, 24:475–483.

61. Bustillos BA, Cocker LT, Coban MA, Weber CA, Bredenberg JM, Boneski PK, Siuda J, Slawek J, Puschmann A, Narendra DP, et al: Structural and Functional Characterization of the Most Frequent Pathogenic PRKN Substitution p.R275W. Cells 2024, 13.

62. Lubbe SJ, Bustos BI, Hu J, Krainc D, Joseph T, Hehir J, Tan M, Zhang W, Escott-Price V, Williams NM, et al: Assessing the relationship between monoallelic PRKN mutations and Parkinson’s risk. Hum Mol Genet 2021, 30:78–86.

63. Castelo Rueda MP, Raftopoulou A, Gogele M, Borsche M, Emmert D, Fuchsberger C, Hantikainen EM, Vukovic V, Klein C, Pramstaller PP, et al: Frequency of Heterozygous Parkin (PRKN) Variants and Penetrance of Parkinson’s Disease Risk Markers in the Population-Based CHRIS Cohort. Front Neurol 2021, 12:706145.

64. Zhu W, Huang X, Yoon E, Bandres-Ciga S, Blauwendraat C, Billingsley KJ, Cade JH, Wu BP, Williams VH, Schindler AB, et al: Heterozygous PRKN mutations are common but do not increase the risk of Parkinson’s disease. Brain 2022, 145:2077–2091.

65. Needham M, Mastaglia FL: Inclusion body myositis: current pathogenetic concepts and diagnostic and therapeutic approaches. Lancet Neurol 2007, 6:620–631.

66. Prasuhn J, Borsche M, Hicks AA, Gogele M, Egger C, Kritzinger C, Pichler I, Castelo-Rueda MP, Langlott L, Kasten M, et al: Task matters - challenging the motor system allows distinguishing unaffected Parkin mutation carriers from mutation-free controls. Parkinsonism Relat Disord 2021, 86:101–104.

67. Hach A, Lohmann, K., Funayama, M., Menon, P.J., Vollstedt, E., Kleinz, T., Yoshino H., Lesage, S., Meier, B., Sue, C.M., Corvol, J.C., Brice, A., Hattori, N., Klein, C., Rakovic, A.: Alternative translation initiation in PRKN delays the onset of Parkinson’s disease and offers a therapeutic target. Annals of Neurology 2026.

68. Mighell TL, Toledano I, Lehner B: SUNi mutagenesis: Scalable and uniform nicking for efficient generation of variant libraries. PLoS One 2023, 18:e0288158.

69. Livesey BJ, Marsh JA: Variant effect predictor correlation with functional assays is reflective of clinical classification performance. Genome Biol 2025, 26:104.

