## Supplementary Table 4 for "A Functional Genetic Atlas of Parkin Resolves Variants of Uncertain Significance and Predicts Parkinson’s Disease Age at Onset"

Supplementary Table 4. Antibodies used in this study

| **Name/Target** | **Species** | **Supplier** | **Catalogue**  **No.** | **Use** | **Concentration/**  **Comment** |
| --- | --- | --- | --- | --- | --- |
| Phospho-ubiquitin (Ser65) | Rabbit (monoclonal) | Cell Signalling Technology | #62803 | FACS (primary antibody) | 1.25 ng/µL |
| Anti-rabbit-Alexa Fluro 647 | Goat | ThermoFisher Scientific | #A21244 | FACS (secondary antibody) | 4 μg/mL |
| Total Parkin (PE-conjugated) | Mouse (monoclonal) | Santa Cruz Biotechnology | sc-32282-PE | FACS | 5 µg/mL |
| Anti-Parkin (PARK2) | Sheep (polyclonal) | MRC PPU Reagents and Services at the University of Dundee | S966C | Immunoblotting (primary antibody) | 1:2000 |
| Anti-Parkin phospho-Ser65 | Rabbit (monoclonal) | Epitomics / Abcam (MJFF collaboration) | - | Immunoblotting (primary antibody) | 1:2000 |
| Anti-Ubiquitin phospho-Ser65 (E2J6T) | Rabbit (monoclonal) | Cell Signalling Technology | 628002S | Immunoblotting (primary antibody) | 1:2000 |
| Anti-Parkin (PRK8) | Mouse (monoclonal) | Santa Cruz Biotechnology | sc-32282 | Immunoblotting (primary antibody) | 1:1000 |
| Anti-GAPDH (6C5) | Mouse (monoclonal) | Santa Cruz Biotechnology | sc-32233 | Immunoblotting (primary antibody) | 1:10,000 |
| Anti-goat | Donkey | LI-COR | 926-68074 | Immunoblotting (secondary antibody) | 1:20,000 |
| Anti-rabbit | Donkey | LI-COR | 926-32213 | Immunoblotting (secondary antibody) | 1:20,000 |
| Anti-mouse | Donkey | LI-COR | 926-32212 | Immunoblotting (secondary antibody) | 1:20,000 |
